# Breaking the bottleneck: self-supervised deep learning framework for fully automated fossil CT segmentation

**DOI:** 10.64898/2026.06.07.730692

**Authors:** Arindam Roy, Poulami Ghosh, Fraser Weston, Ben Hartley, Arianna Salili-James, Sanson T.S. Poon, Susannah C. R. Maidment, Richard J. Butler

**Affiliations:** School of Geography, Earth and Environmental Sciences, University of Birmingham, Birmingham, U.K.; College of Medicine and Health, University of Birmingham, Birmingham, U.K.; Digital, Data and Informatics, Natural History Museum, London, U.K.; AI Lab, Natural History Museum, London, U.K.; Fossil Reptiles, Amphibians and Birds Section, Natural History Museum, London, U.K.

## Abstract

Semantic segmentation of domain-specific imaging data where labelled training examples are scarce and foreground–background contrast is low remains an open challenge in deep learning applied to science. Palaeontological computed tomography (CT) exemplifies this problem: digitally isolating fossilised bone from surrounding rock matrix is labour-intensive (≥100 hrs/dataset), subjective, and often reliant on expensive proprietary software, creating a “segmentation bottleneck” that prevents large-scale and rapid processing of CT data collections. Here we present a self-supervised, end-to-end framework combining SimCLR v1 contrastive pretraining with deterministic pseudo-label generation and U-Net refinement to fully automate fossil CT segmentation without manual annotation. Using 50,626 CT images from the Middle Jurassic Kilmaluag Formation spanning amphibians, reptiles, dinosaurs, and early mammals, the framework achieved a Dice coefficient of 93.66% and IoU of 82.42% on a held-out specimen not seen during training, comparable to the highest Dice and IoU values reported in recent Deep Learning-based fossil CT segmentation studies. Cross-taxon generalisation was validated geometrically on six fully external specimens, achieving sub-voxel mesh agreement with manually thresholded references. By eliminating the annotation requirement that has limited prior deep learning approaches in palaeontology, this framework reduces per-specimen processing from ∼100 person-hours to 6 hrs (one-time UNet training) +1-3 minutes (mesh generation per specimen), an essential first step towards batch processing and analysis of CT data for large-scale comparative and quantitative analyses.

## Main

Since 1984, the progressive integration of X-ray Computed Tomography (CT) into standard palaeontological toolkits has revolutionised the study of fossil anatomy, triggering a methodological shift comparable in impact to the introduction of cladistics or radiometric dating^1,2^. No longer limited to surface exposures or painstaking physical preparation, researchers can now use X-rays to generate non-destructive, three-dimensional (3D) visualisations of internal anatomy with micron to nanometre-level precision (i.e., µCT and SRXTM: Synchrotron X-ray CT). This capability has given rise to the sub-discipline of ‘Virtual Palaeontology’, encompassing the digital acquisition, processing, and analysis of fossil tomographic data^1,2^. Importantly, the value of CT extends far beyond visualisation alone. It now serves as an elementary data source for rigorous quantitative approaches, including Geometric Morphometrics (GM)^3^, Finite Element Analysis (FEA)^4^, and Multibody Dynamics Analysis (MDA)^5^ that are able to shed light on the palaeobiology of extinct organisms by moving beyond the fossilised skeleton to elucidate form and function. Despite these advances, digitally isolating, reconstructing, and annotating small skeletal elements (i.e., the process of ‘segmentation’) remains labour-intensive, shows inter-operator variability, and depending upon anatomical complexity, is extremely time consuming^6–9^. Yu, et al. ^6^, Edie, et al. ^7^ and Knutsen and Konovalov ^9^, provide direct evidence that this time cost often runs into weeks and months. On average, if we take the estimate of 1 hr/slice^9^, even for a small vertebrate dataset of 100-500 slices (many datasets are frequently larger), the calculated minimum time-cost ceiling would conservatively be ≥100 person-hours.

Segmentation workflows often require expensive proprietary software and substantial domain-specific anatomical expertise. This has created a profound ‘segmentation bottleneck,’ whereby CT data are generated faster than they can be processed within realistic academic timeframes, severely constraining large-scale palaeontological research and limiting the scientific potential of existing CT collections. Furthermore, the time cost that is incurred during manual segmentation could be better budgeted to downstream anatomical, phylogenetic, and functional analyses. This challenge is also an open problem in machine learning (ML) and deep learning (DL): fossil CT data presents extreme foreground–background contrast ambiguity, severe class imbalance due to incomplete preservation, and morphological diversity that exceeds the assumptions of standard segmentation architectures trained on biomedical or natural photographic datasets (e.g., ImageNet).

A foundational technique in virtual palaeontology is global thresholding^2^. Also referred to as iso-surfacing, this method assumes that fossilised material and the enclosing matrix occupy distinct, non-overlapping ranges of radiodensity in images, expressed as attenuation coefficients^10^. In practice, however, the bimodal histogram required for clean thresholding is the exception. Fossilisation is a complex geochemical process: mineral replacement of biological tissues frequently produces fossils with radiodensity values nearly identical to the enclosing matrix. Calcareous shells in limestone, for example, exhibit negligible density contrast, while vertebrate fossils in iron-rich concretions suffer from beam hardening — a CT artefact in which preferential absorption of low-energy X-rays causes artificial density gradients that confound intensity-based separation. The result is segmentation errors including matrix bleed, loss of thin structures such as turbinates or trabeculae, and artificial fusion of adjacent skeletal elements^11–13^. As a result, researchers were forced to adopt local adaptive thresholding (or manual painting approaches).

In response, the software industry introduced semi-automated segmentation solutions. Commercial platforms such as Avizo^14^, Mimics^15^, Volume Graphics StudioMAX^16^, and Dragonfly^17^ became standard tools in digital palaeontology. These packages implemented algorithms including region growing/shrinking^18^, watershed segmentation^19^, and slice interpolation^20^, allowing users to define seed regions or segment every n^th^ slice while the software interpolated intervening volumes. More advanced tools, such as Dragonfly’s Segmentation Wizard^21^ and the Trainable Weka Segmentation^22^ plugin in ImageJ, employ classical Machine Learning classifiers, most commonly Random Forests^23^ or Support Vector Machines^24^, operating on local texture features. In these workflows, the user manually annotates a subset of voxels to define classes such as bone, matrix, or air, and the classifier predicts labels for the remainder of the volume based on voxel intensity and local texture statistics. Although these approaches represented a substantive advance over fully manual segmentation, several limitations prevent effective scaling^6,8^. First, they require a persistent human-in-the-loop and remain fundamentally interactive rather than automated. Domain experts must generate training data for each specimen, rendering these workflows impractical for large-scale segmentation of CT datasets. Second, the resulting classifiers are typically specimen-specific, overfitted to the contrast, noise characteristics, and morphology of individual scans, and rarely transferable without retraining. As a result, training effort is effectively discarded after each session, leading to extensive duplication of labour across the field. Third, reliance on proprietary software undermines reproducibility and accessibility. Segmentation workflows implemented in commercial platforms often function as black boxes, with algorithmic details, feature selection, and parameterisation obscured. While Dragonfly offers a FreeD licence for qualifying academic users, this programme excludes government institutions, non-profit organisations, and researchers in several major research territories including Japan, South Korea, Singapore, and Mainland China including Hong Kong and Macau ^25,26^. Avizo, Mimics, and VGStudioMAX have no equivalent free academic licensing programmes. Furthermore, the ML/DL modules within these commercial platforms operate as supervised, specimen-specific workflows requiring per-specimen manual annotation, architecturally distinct from the annotation-free, self-supervised framework presented here. For a complete comparison of features across different segmentation software and this study, please refer to **Supplementary Information Table, S1**.

Given the substantial time investment, labour demands, specialised expertise, and software costs involved, the segmentation bottleneck represents an ideal target for automation using artificial intelligence (AI)^6,8,9^.

Since 2020, the palaeontological community has increasingly explored DL- approaches, particularly Convolutional Neural Networks (CNNs)^8^, building on their proven success in biomedical image analysis, where DL has revolutionised image segmentation, achieving near-human performance in delineating soft tissues, tumours, and organs. Early fossil CT segmentation workflows understandably drew on established approaches, including classical machine-learning methods or relatively shallow CNN architectures^27–29^. These methods often relied on limited sets of handcrafted grayscale, texture, or edge-based features, which proved effective under well-controlled contrast conditions but were less able to capture broader morphological context across complex fossil geometries. Subsequent adoption of encoder-decoder architectures such as U-Net marked a significant methodological advance by enabling end-to-end feature learning. However, these early applications were typically focused on narrowly defined anatomical regions^12,30^ or single taxonomic groups including foraminifera^31^, bivalves^7^, vertebrate microfossils^28^, and dinosaur embryos^6^. As a result, the demonstrated generalisability of these models across phylogenetic breadth, ontogenetic stages, and diverse taphonomic regimes remains limited. Reported high Dice-Sørensen coefficient (DICE) or Intersection-over-Union (IoU, also known as Jaccard Index) scores in this context often derive from small or highly curated training datasets, sometimes comprising fewer than ten specimens or a limited number of annotated slices. While such results highlight the promise of DL-based segmentation, they may also underestimate the impact of morphological and preservational variability, thereby inflating apparent performance.

More recent studies employing advanced architectures, including EfficientNet + U-Net^7^ and ResNet-backboned U-Net^32^, U-Net++^32^, 2.5D CNNs^32^, and hybrid Vision Transformer- generative adversarial network (ViT-GAN)^33^ pipelines, have achieved notable improvements in boundary delineation and annotation efficiency. Techniques such as few-shot learning and GAN based augmentation provide valuable strategies for mitigating limited sample sizes. However, they may also reinforce dataset-specific characteristics when synthetic volumes do not fully capture the heterogeneity of fossil preservation or matrix composition encountered in broader applications. In addition, reports of early accuracy plateaux following minimal annotation or training effort suggest that some models may overfit to narrowly defined datasets and tolerate relatively high false-positive or false-negative rates when applied beyond the original specimens or scanning conditions^6,9,31^.

To address these persistent challenges, we introduce a self-supervised, end-to-end segmentation framework that effectively bypasses the need for initial manual annotation. Our framework utilises SimCLR v1 (Simple Framework for Contrastive Learning of Visual Representations ver.1)^34^ as a pre-training phase, where it learns high-level feature representations directly from raw, unlabelled CT slices through stochastic augmentations. Concomitantly, we implemented a suite of rule-based, deterministic image processing algorithms to generate a standardised preliminary set of ground truth masks. These two independent information streams are integrated through a knowledge fusion step (**Fig. 1**), i.e., the integration of heterogeneous information sources into a unified model representation, to train a modified U-Net architecture which proceeds to refine coarse mask predictions into high-precision overlays that closely match the anatomical regions of interest, even in datasets with low contrast differences. Unlike traditional workflows that discard training effort after each specimen, this framework was implemented on a dataset of 50,626 CT image slices. This dataset, derived from vertebrate fossils comprising amphibians, reptiles, dinosaurs and early mammals, represents a five-fold increase over the largest training set previously employed in palaeontological CT segmentation (7,986 slices)^6^, and is orders of magnitude larger than the 5-18 manually annotated slices used in most recent fossil CT segmentation studies^7,9,30^.

**Fig 1.**
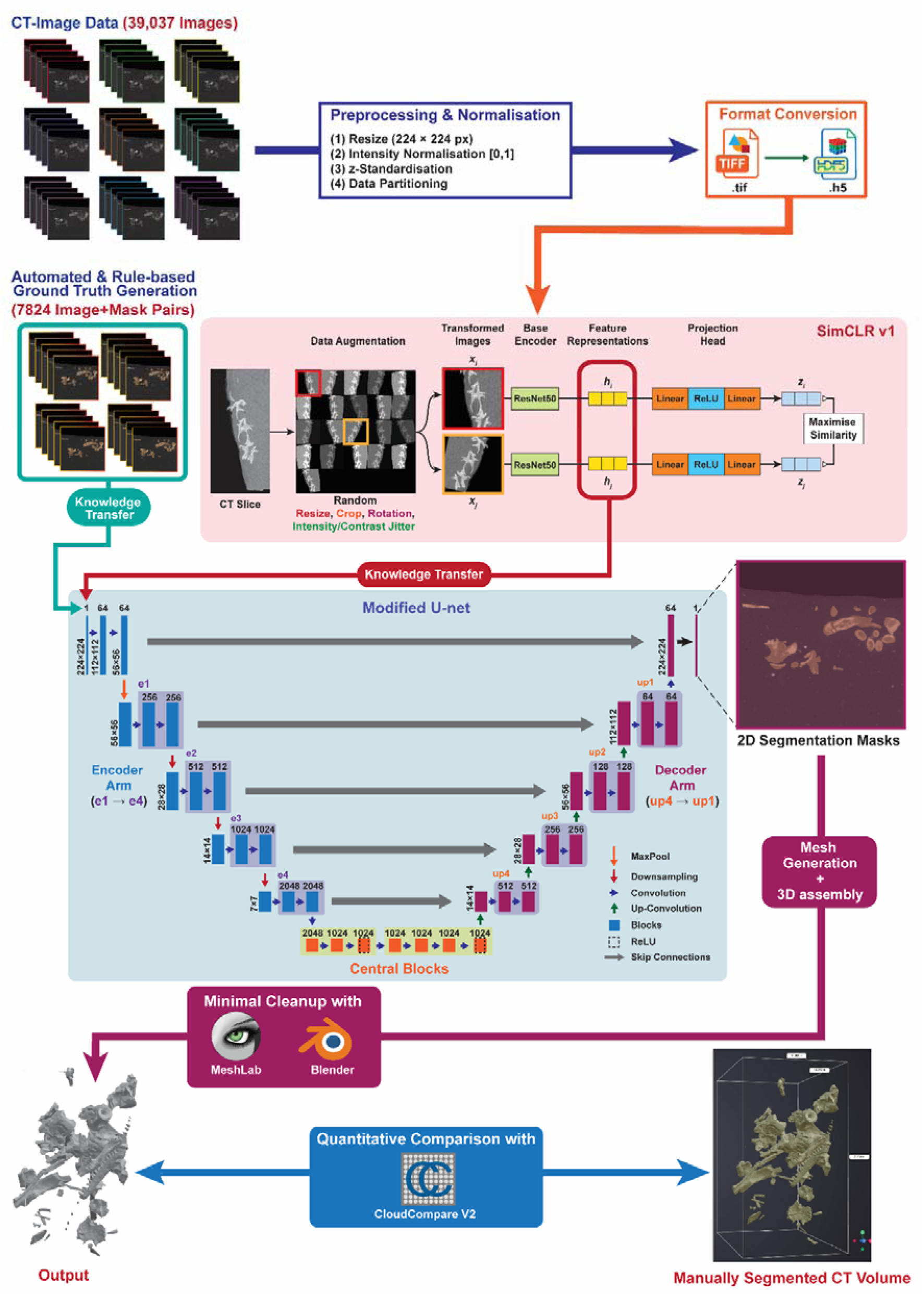
The core workflow for AI based automated segmentation of CT data proposed in this study. The workflow has three components: the raw CT data is first subjected to preprocessing and passed on to SimCLR. In parallel coarse mask-image pairs are generated using a sequence of deterministic image transformation. The feature representations from SimCLR and coarse mask-image pairs are passed on through separate knowledge transfer steps to U-Net which generates the fine masks for the 2D slices. These masks are then stitched together to generate the final mesh which is visualised compared against the gold standard manually segmented (binary thresholded) using MeshLab, Blender and CloudCompare v2 respectively.

The unprecedented scale and taxonomic breadth of this dataset are critical for establishing a reproducible alternative to human-in-the-loop segmentation. We specifically selected the fossils from the Kilmaluag Formation, Scotland^35^, as ‘the challenge dataset’ for this framework due to the site’s exceptional scientific value (officially designated as a Site of Special Scientific Interest by NatureScot) as one of two localities worldwide that yield abundant and relatively complete skeletons of small land-dwelling animals from the Middle Jurassic Period. These fossils are typically preserved in consolidated carbonate, where the mineral replacement of biological tissues often results in a radiodensity nearly identical to the surrounding matrix, rendering global thresholding somewhat unproductive in many cases. Our methodological contribution is the demonstration that, in a domain where supervised training data is structurally scarce, combining contrastive self-supervision on domain-specific imagery with deterministic pseudo-labels provides a sufficient signal for high-precision semantic segmentation without any manual annotation. Our application contribution is the first fully automated, end-to-end CT segmentation pipeline for palaeontology, validated across multiple vertebrate taxa, robust to noise, mineral-density contrasts, and scan-acquisition artefacts within a broadly similar taphonomic regime (**Fig. 1**).

## Results

### Contrastive pre-training encodes fossil-matrix structure without labels

The SimCLR v1 self-supervised pre-training, conducted on 39,037 unlabelled CT slices drawn from a taxonomically diverse corpus of 50,626 images yielded robust, domain-specific feature representations that reliably discriminate fossilised bone from surrounding geological matrix without requiring manual annotation. Over 250 training epochs, contrastive accuracy converged to 93.89% on the training set and 93.66% on the held-out validation set (**Fig. 2b**), with corresponding normalised temperature cross entropy (NT-Xent) losses of 0.2036 and 0.2312 (**Fig. 2a**). The narrow gap between training and validation metrics indicates effective generalisation rather than memorisation. Both loss curves descend steeply during the first approximately 50 epochs before plateauing (**Fig. 2a**), while accuracy curves rise in parallel with no divergence (**Fig. 2b**), confirming stable convergence without overfitting.

**Fig 2.**
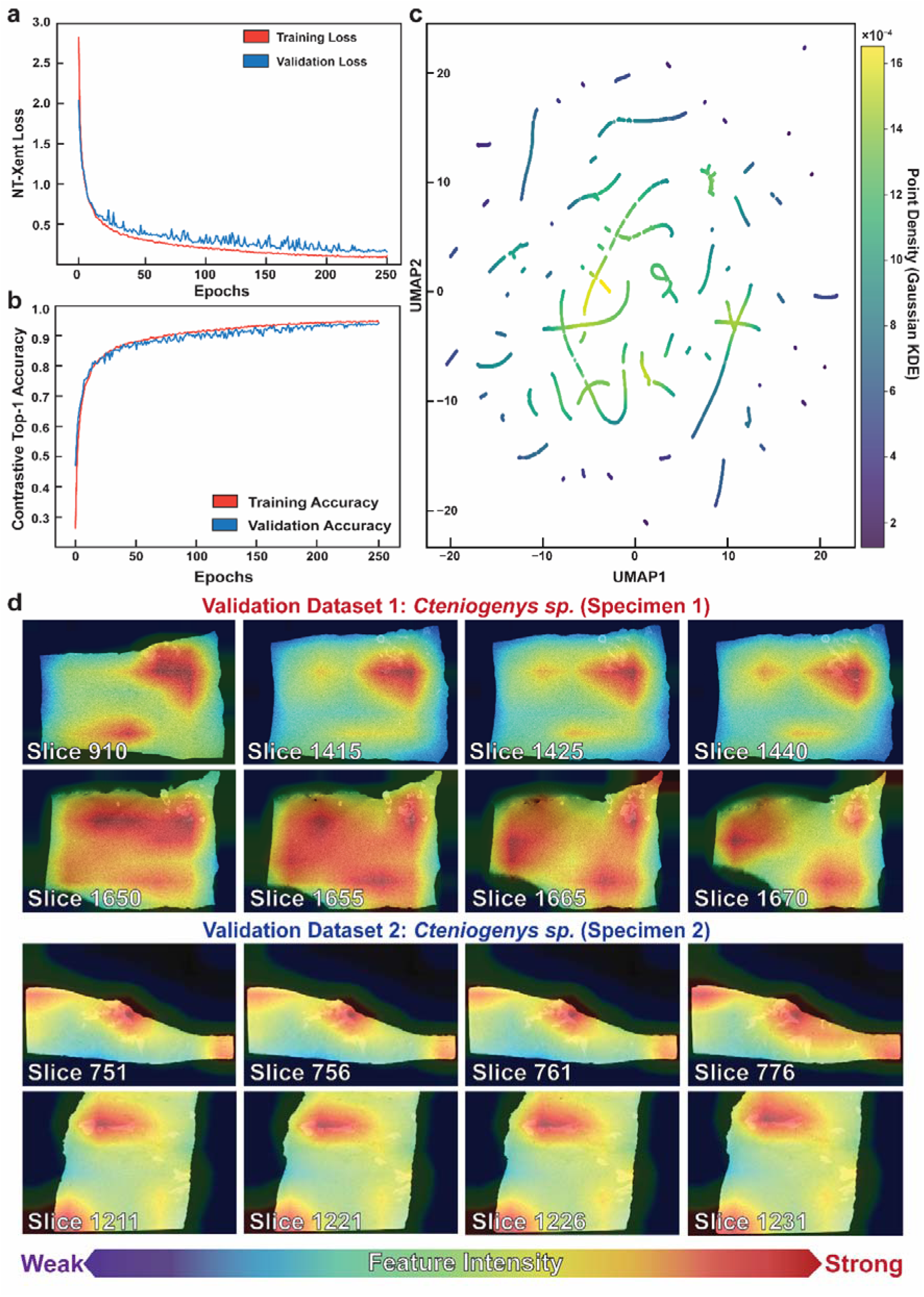
Metrics from SimCLR Training and Validation. Training and validation accuracy (a) and Training and validation loss (b) metrics for SimCLR, UMAP plot for features in validation (c) and visual sanity check for two datasets from Cteniogenys sp. (Choristodera, Specimen1: MorphoSource 000521279, Specimen 2: MorphoSource 000680995) as validation data using Grad-CAM (d): strong activation over skeletal material confirms that the encoder captures biologically meaningful bone-matrix structure.

To verify that the learned representations encode biologically meaningful structure rather than radiographic noise, we applied two complementary post-hoc validation techniques. Uniform Manifold Approximation Project (UMAP) dimensionality reduction^36^ was applied to the 2048-dimensional feature vectors extracted by the trained ResNet-50 encoder^37^, projecting them into two-dimensional space (**Fig. 2c**). The projection is overlaid with a Gaussian Kernel Density Estimate (KDE), mapping point density from dark purple (sparse) through teal and green (moderate) to yellow (highest-density cores). Crucially, no class labels are used; all visible structure has emerged entirely from the contrastive objective. This emergent clustering is characteristic of well-trained contrastive encoders^34,38^ and serves as a standard quality check for self-supervised representations.

Gradient-weighted Class Activation Mapping (Grad-CAM) heatmaps^39^ provided spatially explicit, slice-level confirmation of the UMAP findings. Activation maps, spanning two separate datasets of *Cteniogenys* sp. sampled diverse anatomical cross-sections (**Fig. 2d**). Heatmaps consistently localise strong activation over fossilised skeletal material across both specimens, from dense bone cross-sections to peripheral slices containing smaller fragments. Activation intensity tracks bone spatial extent and density, while matrix regions with elevated radiodensity due to mineralogical inclusions or artefacts do not elicit strong responses, confirming the encoder distinguishes genuine bone texture from superficially similar background signals.

Together, the structured multi-cluster organisation in the UMAP feature space (**Fig. 2c**) and the spatially precise bone localisation in the Grad-CAM heatmaps (**Fig. 2d**) confirm that SimCLR v1 pre-training has yielded a feature representation that partitions CT data along axes directly relevant to downstream segmentation: bone versus matrix discrimination, graded bone density representation, and radiographic artefact rejection.

### Rule-based masking generates reproducible bone-location priors at scale

A critical step enabling end-to-end automation is the parallel generation of standardised ground truth masks through a fully deterministic, rule-based image processing pipeline. Rather than relying on human annotators to manually delineate bone in each CT slice, we developed a sequence of classical image processing operations with fixed parameters, applied identically across the entire test partition (3 datasets, 7,824 images) to produce coarse but consistent binary masks of fossilised bone (**Fig. 3c**, middle column). These masks served not as final segmentation outputs, but as reproducible spatial guidance for the subsequent U-Net training through the knowledge fusion strategy (**Fig. 1**).

**Fig 3.**
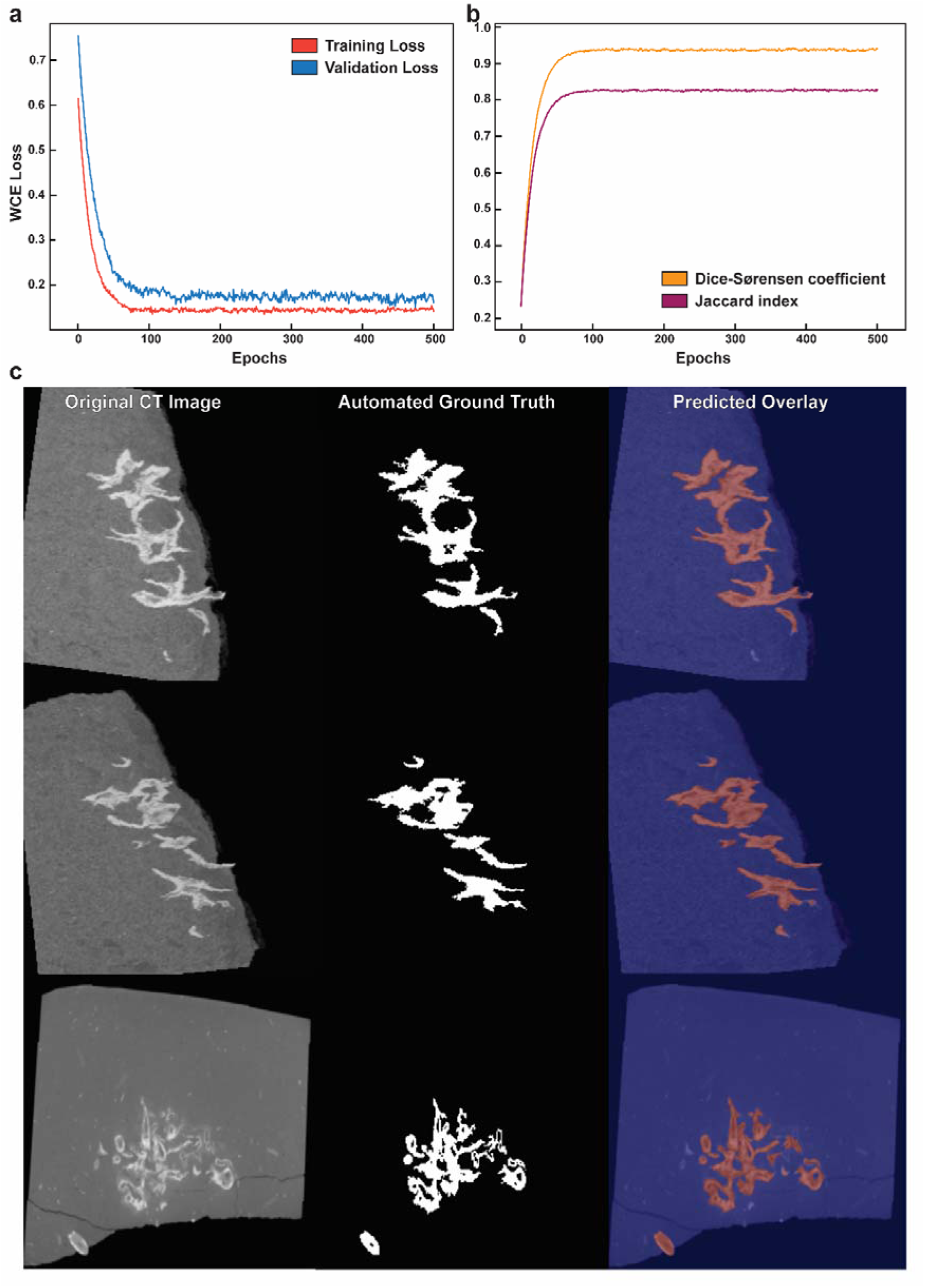
Training performance metrics for U-Net. Training and test loss (a), Dice and IoU metrics evaluated on the held-out test specimen (b). Comparison of the original CT image, rule-based deterministic masks, and U-Net generated fine masks (c). The close tracking of training and test loss in (a) and the stable Dice and IoU values in (b) indicate that the U-Net generalises to the held-out specimen without overfitting; in (c), the U-Net masks (right) recover continuous, fine-scale bone margins from the coarse rule-based masks (middle) while suppressing surrounding matrix.

The output is illustrated in **Fig. 3c**, where three representative CT slices spanning different specimens and anatomical configurations are shown alongside their corresponding automated masks. The left column presents the original CT images, revealing the characteristic challenge of the Kilmaluag Formation material: fossilised bone and dolomitic matrix share similar grey-level intensities. The middle column presents the automated ground truth masks. These masks capture the gross morphology of the skeletal material in each slice, delineating the major bone cross-sections and preserving the overall spatial arrangement of anatomical elements. In the upper two rows, where relatively dense and well-defined bone fragments are present, the masks provide a faithful outline capturing both large contiguous bone masses and smaller isolated fragments. In the bottom row, where the bone is more fragmented and scattered with lower contrast, the algorithm still identifies the principal skeletal elements, though some smaller or lower-contrast fragments are either missed or incompletely captured.

This variation reflects an inherent limitation of the rule-based approach: the masks are simplified outlines that approximate bone extent rather than precisely delineating fossil-matrix boundaries. At ambiguous interfaces where taphonomic density convergence^40^ produces near-identical attenuation coefficients between bone and rock, the algorithm may over-segment (incorporating a matrix halo) or under-segment (failing to capture thin or poorly mineralised margins). The masks are explicitly intended as coarse spatial priors; their role is to provide U-Net with approximate spatial constraints indicating where bone is likely located, while the SimCLR-derived feature representations supply the discriminative textural information needed to resolve ambiguous boundaries. The predicted overlays in **Fig. 3c** (right column) demonstrate substantially refined boundaries compared to the coarse automated masks. All pipeline parameters were determined empirically on the training partition and held fixed thereafter. Given the same input CT slice, the algorithm always produces the identical mask, ensuring full deterministic reproducibility.

### Knowledge fusion yields high-precision 3D bone reconstructions

The knowledge fusion stage combined the two independent information streams, domain-specific feature representations from SimCLR v1 pre-training and standardised spatial guidance from the rule-based automated masks, into a single segmentation model. A modified U-Net architecture^41^ initialised with SimCLR-trained ResNet-50 encoder weights refined the coarse mask-image pairs into high-precision segmentation overlays. Using a composite Weighted Cross-Entropy and Dice loss function^42^ to address the pronounced class imbalance between bone and matrix voxels, training and test losses converged within 50 epochs and plateaued without divergence (Fig. 3a). On the held-out test dataset, the model achieved a Dice coefficient of 0.9366 and IoU of 0.8242 at the best checkpoint (Fig. 3b), confirming generalisation beyond the training data.

The qualitative improvement over the coarse automated masks is directly visible in Fig. 3c. Where relatively well-defined bone cross-sections are present (upper two rows), the U-Net predicted overlays closely follow anatomical boundaries with visibly sharper edges, respecting fine morphological detail including narrow bone processes and thin cortical margins that the rule-based algorithm captured only approximately. Where bone is more fragmented and lower in contrast (bottom row), the model continues to identify the principal skeletal elements while reducing false positive inclusions evident in the corresponding automated mask.

These 2D segmentation masks were stacked to reconstruct volumetric 3D meshes preserving the shape and spacing of the original CT scans. Six external fossil CT datasets, comprising stem salamanders (’Salamander A’ Specimens 1 and 2 and *Marmorerpeton wakei* Specimens 1 and 2), an early mammal (Mammaliaformes indet.), and an undescribed docodont mammaliaform, were segmented through region growing, Otsu-based intensity filtering, and morphological cleanup, followed by surface extraction using the marching cubes algorithm. The resulting watertight triangulated meshes required no manual post-processing and are suitable for downstream geometric morphometric, finite element, and 3D printing applications.

The total one-time computational investment was approximately 43–44 hours (SimCLR pre-training: ∼37–38 hrs; U-Net training: ∼6 hrs 10 min on a single NVIDIA A100 GPU, peak memory ∼3.26 GB), after which the model segments new CT datasets at inference without retraining, producing per-specimen 3D meshes in 1 to 3 minutes. Mesh quality metrics for all six specimens are reported in Table 1. Five of six specimens achieved mean Dice scores of approximately 0.84 and IoU of approximately 0.73, with Standard Tessellation Language (STL) meshes comprising 5–8.4 million vertices and 9.9–16.7 million faces. Mammaliaformes indet. shows lower values (Dice 0.743, IoU 0.596) and a substantially smaller mesh (2.0 million vertices, 4.0 million faces), attributable to the presence of multiple minuscule bone fragments and prominent ring artefacts in the source CT data.

**Table 1.** Segmentation quality and mesh output for six external fossil specimens. Mean Dice-Sørensen coefficient and Intersection-over-Union (IoU) are derived from the underlying 2D segmentation masks; STL vertex and face counts reflect surface mesh complexity. Mask shape is reported as Z × Y × X slices. Lower values for Mammaliaformes indet. are attributable to ring artefacts in the source CT data.

| Specimen | Mask shape ($Z \times Y \times X$) | STL vertices | STL faces | Mean Dice | Mean IoU | Mesh Generation Time (hr:min:sec) |
| --- | --- | --- | --- | --- | --- | --- |
| Salamander A (Specimen 1; MorphoSource: 00084381) | $2057 \times 1192 \times 788$ | 7,664,089 | 15,205,856 | 0.838 | 0.723 | 00:03:05 |
| Salamander A (Specimen 2; rescan of 000071513) | $1496 \times 672 \times 1198$ | 5,872,767 | 11,729,270 | 0.843 | 0.730 | 00:02:42 |
| *Marmorerpeton wakei* (Specimen 1; MorphoSource: 000700518) | $1591 \times 1322 \times 991$ | 5,002,740 | 9,989,358 | 0.838 | 0.723 | 00:02:08 |
| *Marmorerpeton wakei* (Specimen 2; MorphoSource: 000700519) | $1901 \times 1114 \times 1022$ | 8,385,631 | 16,719,920 | 0.843 | 0.730 | 00:03:21 |
| *Mammaliaformes* indet. (MorphoSource: 000693738) | $1955 \times 970 \times 1163$ | 2,010,240 | 3,984,930 | 0.743 | 0.596 | 00:01:00 |
| Docodonta (undescribed; MorphoSource: 000721884) | $1906 \times 1257 \times 1054$ | 7,789,899 | 15,560,350 | 0.840 | 0.728 | 00:03:18 |

### Our AI-framework generated meshes achieve sub-voxel agreement with manual thresholding

To assess the fidelity of the AI-generated meshes against an established palaeontological workflow, the six specimens segmented by the SimCLR+U-Net pipeline were independently segmented using binary thresholding in Avizo at multiple threshold levels (**Supplementary Information**, **Fig. S1**). For each specimen, both meshes were scaled to a common 1:1 coordinate space using their respective voxel sizes prior to registration, ensuring dimensional equivalence without relying on algorithmic scale adjustment. The resulting mesh pairs were then compared through a three-phase geometric registration protocol in CloudCompare v2.

First, Point Pair Registration (PPR)^43^ established a common coordinate system through manual landmarking, yielding coarse alignments with RMS errors ranging from 0.081 mm to 0.363 mm across the assemblage. Second, Iterative Closest Point (ICP) refinement^44^ minimised nearest-neighbour distances through rigid transformation (rotation and translation only), with scale fixed at 1.0 throughout to confirm that the pre-registered meshes are dimensionally equivalent and to prevent the algorithm from artificially warping either mesh toward noise clusters. Third, Cloud-to-Cloud (C2C)^45^ signed distance analysis quantified per-point deviations between the full automated point cloud and the manual reference surface, yielding mean distance, standard deviation, and Hausdorff (maximum) distance statistics.

For Salamander A (Specimen 1), Salamander A (Specimen 2) and Docodonta (undescribed), ICP overlap was set to 100%, meaning all points were used and the resulting RMS captures full geometric divergence. For Mammaliaformes indet., *Marmorerpeton wakei* (Specimens 1 and 2), overlap was set to 80%, computing correspondences on only the best-matching 80% of points and discarding the worst 20% as outliers during each iteration. This trimmed setting was applied to specimens exhibiting significant non-overlapping geometry: multi-element vertebral complexes and fragmented specimens with prominent ring artefacts, where outlier fragments would otherwise bias the rigid alignment of core anatomy. The C2C signed distances in the third phase were computed on the full, untrimmed point clouds for all six specimens regardless of the ICP overlap setting (**Fig. 4**).

**Fig 4.**
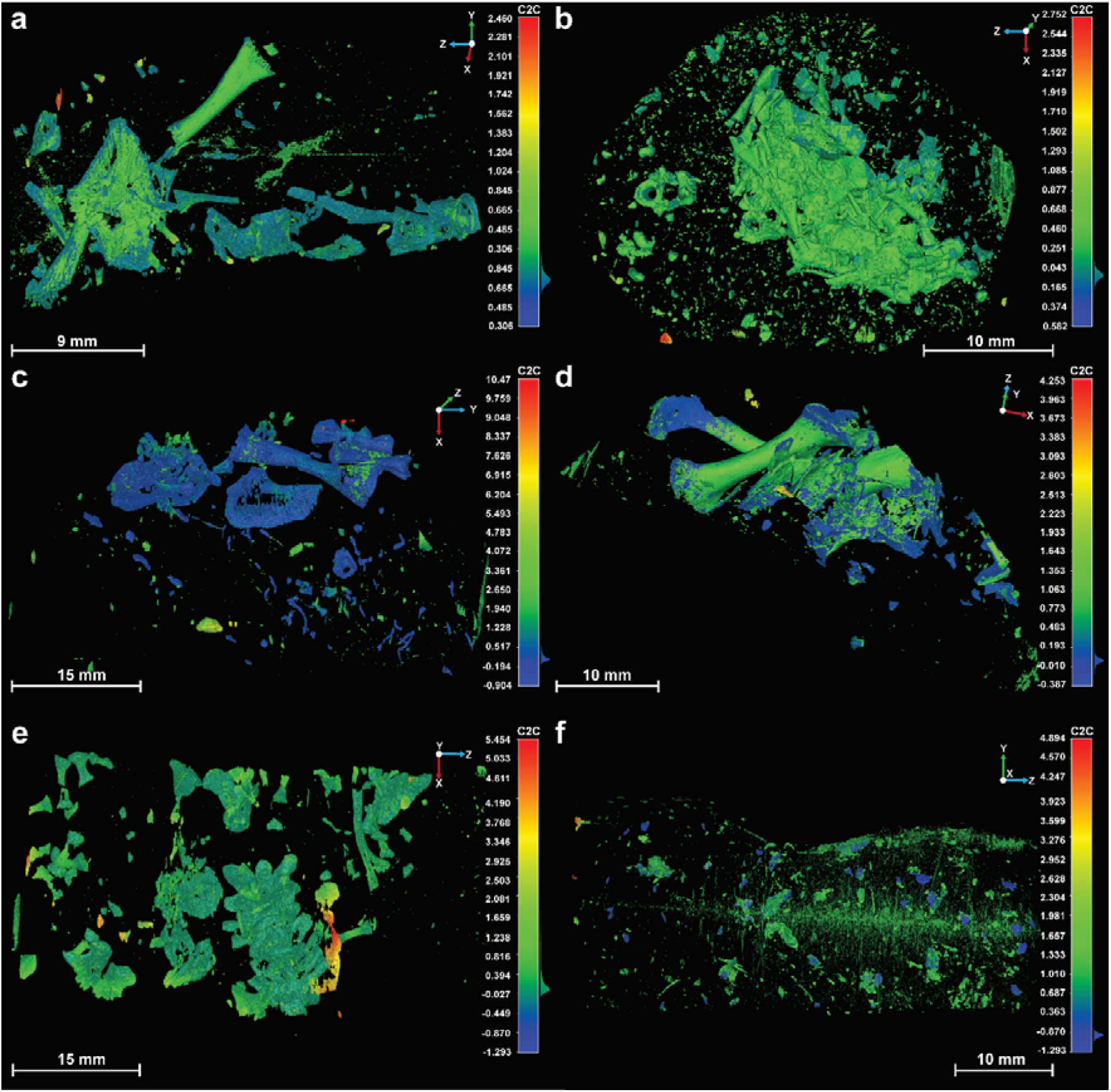
Results of Cloud Compare investigating the similarity between AI generated and manually segmented meshes. (a) Salamander A (Specimen 1; MorphoSource: 00084381), (b) Salamander A (Specimen 2; high-resolution rescan of MorphoSource 000071513), (c) Marmorerpeton wakei (Specimen 1; MorphoSource: 000700518), (d) Docodonta (undescribed; MorphoSource: 000721884), (e) Marmorerpeton wakei (Specimen 2; MorphoSource: 000700519), (f) Mammaliaformes indet. (MorphoSource: 000693738).

The registration results reveal a clear three-tier performance hierarchy across the assemblage (**Table 2**). Tier 1 (sub-voxel agreement): Salamander A (Specimen 2) (C2C mean = −0.017 mm, 0.75 voxels) and Docodonta (undescribed) (C2C mean = 0.016 mm, 0.63 voxels) achieve sub-voxel agreement between the AI-generated and manually thresholded meshes, meaning the two segmentation methods resolve essentially identical geometry at the resolution limit of the CT data. The C2C mean for Salamander A (Specimen 2) is negative, indicating that the SimCLR+U-Net segmentation produces slightly tighter bone boundaries than binary thresholding. The C2C colour maps for these specimens (**Fig. 4b, d**) show predominantly green surfaces with tight colour distributions.

**Table 2.**
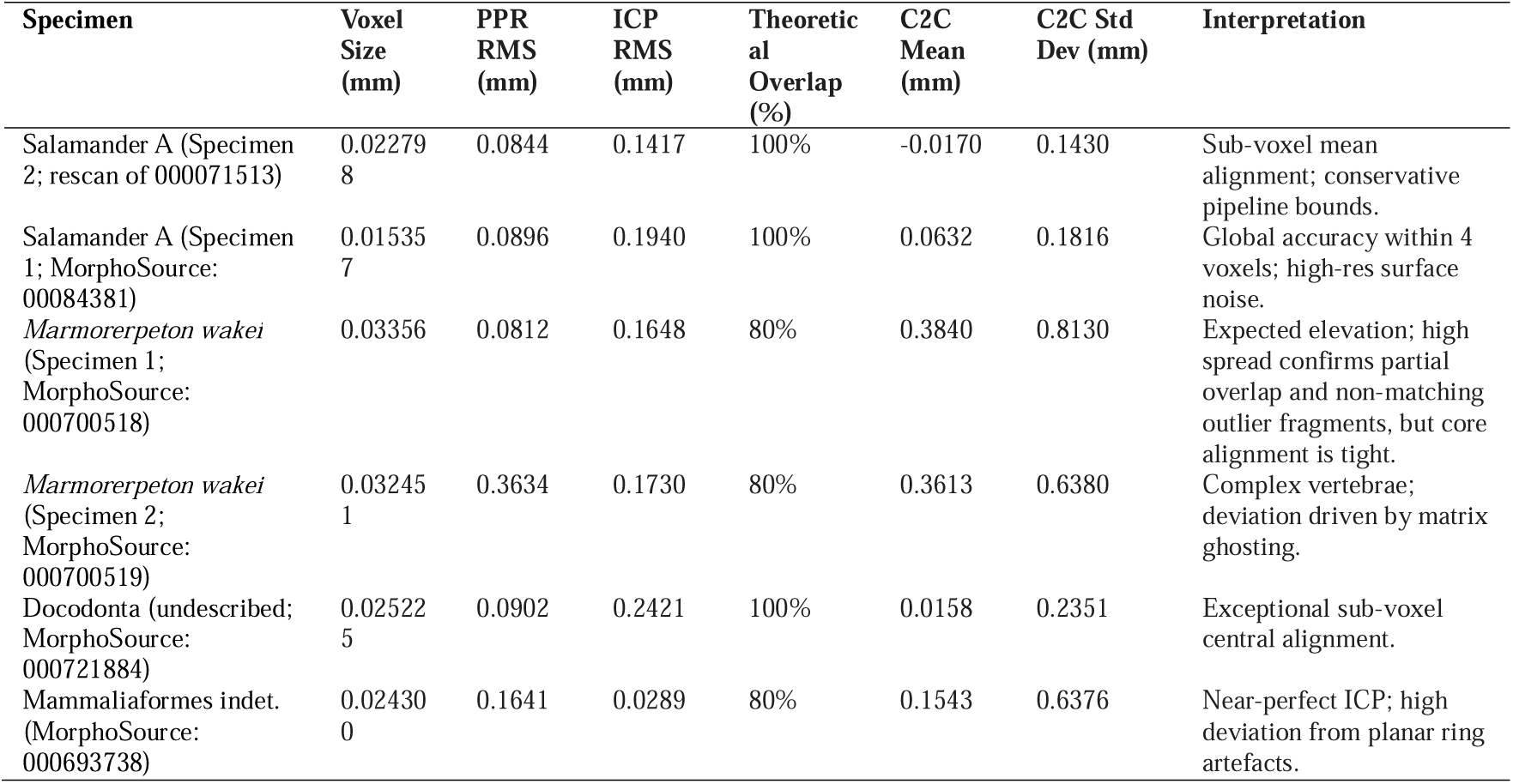
Geometric registration statistics between AI-generated and manually thresholded meshes for six specimens. Point-pair Registration (PPR) and Iterative Closest Point (ICP)-Root Mean Squared (RMS) errors quantify coarse and fine alignment; Cloud-to-Cloud (C2C) mean distance and standard deviation are computed on full, untrimmed point clouds. All distances in mm. Theoretical overlap indicates the proportion of points used during ICP refinement.

Salamander A (Specimen 1) (C2C mean = 0.063 mm, 4.11 voxels) and Mammaliaformes indet. (C2C mean = 0.154 mm, 6.35 voxels) constitute Tier 2 (low absolute deviation with scan-dependent voxel ratios). For Salamander A (Specimen 1), the relatively fine voxel resolution of 0.0154 mm means the elevated voxel ratio reflects the higher resolving power of the scan rather than poor absolute agreement; the absolute C2C mean of 0.063 mm is within the range expected from surface noise at this resolution. Mammaliaformes indet. presents a distinctive pattern: the best ICP RMS in the assemblage (0.029 mm, 1.19 voxels at 80% overlap), indicating near-perfect alignment of core bone mass, but elevated C2C spread (std = 0.638 mm, 26.24 voxels). This is attributable to planar ring artefacts visible as horizontal banding in the C2C colour map (**Fig. 4f**), present in the Avizo binary threshold mesh but absent from the AI-generated output.

*Marmorerpeton wakei* (Specimen 1) (C2C mean = 0.384 mm, 11.44 voxels) and *Marmorerpeton wakei* (Specimen 2) (C2C mean = 0.361 mm, 11.13 voxels) constitute Tier 3 (elevated metrics driven by non-overlapping peripheral geometry). Both specimens are multi-element vertebral complexes where scattered fragments are captured by one method but not the other, inflating distance statistics. Where the two methods share anatomy, alignment is tight: *Marmorerpeton wakei* (Specimen 1) has the best PPR coarse alignment (0.081 mm, 2.42 voxels), with a near-identity transformation matrix (off-diagonal rotation terms on the order of 0.001 to 0.006, translations below 0.17 mm), and the subsequent ICP correction is even smaller (sub-micron rotations, translations below 0.02 mm). The C2C colour maps for these specimens (**Fig. 4c, e**) show blue scatter from unmatched fragments surrounding the green-aligned vertebral chain, confirming that the elevated statistics are driven by peripheral non-overlapping geometry.

## Discussion

This study presents a fully automated, self-supervised segmentation framework that removes the need for human annotation in palaeontological CT workflows and achieves sub-voxel geometric agreement with established manual methods for well-preserved specimens. For specimens affected by ring artefacts or fragmentation, the pipeline correctly excludes reconstruction noise that manual thresholding retains. The SimCLR+U-Net pipeline, trained on 39,037 unlabelled CT slices spanning amphibians, reptiles, dinosaurs, and early mammals, attained a Dice coefficient of 0.9366 and IoU of 0.8242 on a held-out test specimen not seen during training, comparable to the highest Dice and IoU values reported in recent DL-based fossil CT segmentation studies^6,7,9,30–32,46^. External geometric comparison against binary thresholding in Avizo yielded C2C means of 0.63 to 0.75 voxels for the best-performing specimens at a fixed scale of 1.0.

The adoption of SimCLR v1 as a self-supervised pre-training strategy addresses the most persistent obstacle to applying deep learning in palaeontological CT: the need for large volumes of expert-annotated training labels. In standard workflows, specialists manually trace bone boundaries on each training slice, and this effort is discarded after each specimen and repeated from scratch for new material. The contrastive objective used here learns directly from 39,037 unlabelled slices drawn from a corpus of 50,626 images, extracting feature representations that capture fossil textures, taphonomic noise signatures, and beam hardening patterns without any supervisory signal. This corpus substantially exceeds all previously published fossil CT training sets in both scale and taxonomic breadth. The representations learned on palaeontological material are expected to outperform generic ImageNet-pretrained weights^47^ for fossil-matrix discrimination, because ImageNet features are optimised for natural-image statistics (colour, texture, object shape) that have little in common with the greyscale, low-contrast characteristics of µCT data. The SimCLR augmentation strategy was designed to match the structural characteristics of palaeontological CT data. Geometric transforms (random rotation, flipping, and scale jitter) enforce invariance to arbitrary specimen orientation within the matrix. Intensity and blur augmentations enforce invariance to the scanner- and matrix-chemistry-dependent variation in CT attenuation values characteristic of multi-institutional fossil datasets. These are precisely the sources of variance that cause models pretrained on biomedical or natural-image corpora to fail on fossil material To our knowledge, this represents the first application of contrastive self-supervised learning to palaeontological CT segmentation. The contrastive objective also acts as an implicit regulariser: by requiring the encoder to map different augmentations of the same slice (random cropping, rotation, intensity jitter, Gaussian blur) to nearby positions in latent space, SimCLR builds invariance to exactly the sources of variability that cause supervised models to fail on small palaeontological datasets, including scan noise, orientation differences, and contrast variation between specimens.

The knowledge fusion strategy, in which coarse rule-based masks provide spatial constraints and SimCLR-derived features provide discriminative power, marks a shift from prior palaeontological deep learning workflows that depend on specimen-specific, manually created ground truth. In those approaches, training labels are produced by hand for each new specimen and discarded after the model is trained, duplicating effort across the field. The deterministic masking pipeline developed here generates reproducible spatial priors using fixed parameters, while the SimCLR encoder supplies the textural discrimination needed to resolve ambiguous fossil-matrix boundaries that the rule-based algorithm cannot handle. The U-Net learns to correct the over- and under-segmentation of the coarse masks through this dual-stream transfer, producing refined boundaries that are more accurate than either input alone. Once the framework is trained, it segments new specimens without any additional annotation, and the accumulated training signal is preserved and reusable rather than discarded after each session.

The geometric validation provides quantitative evidence that the pipeline preserves the true physical dimensions of biological material. We classify the six external specimens into three tiers based on C2C distance statistics: Tier 1 (sub-voxel agreement, C2C mean <1 voxel), where the two segmentation methods resolve essentially identical geometry at the resolution limit of the CT data; Tier 2 (low absolute deviation, C2C mean 1–7 voxels), where elevated voxel ratios reflect scan resolution or artefact contamination rather than segmentation error; and Tier 3 (elevated metrics, C2C mean >10 voxels), where distance statistics are inflated by non-overlapping peripheral fragments rather than poor accuracy of the shared anatomy. Salamander A (Specimen 2) (C2C mean = 0.75 voxels) and Docodonta (undescribed) (C2C mean = 0.63 voxels) fall within Tier 1, achieving sub-voxel agreement between my AI-pipeline generated and manually thresholded meshes, with both meshes pre-scaled to a common 1:1 coordinate space using their respective voxel sizes and ICP scale held at 1.0. The strongest evidence for improvement over binary thresholding comes from how the pipeline handles artefacts. The Mammaliaformes indet. specimen (Tier 2) achieves the best ICP RMS in the assemblage (1.19 voxels for core bone mass at 80% overlap), yet its C2C standard deviation is inflated to 26.24 voxels by planar ring artefacts. These artefacts are present in the Avizo threshold mesh but are correctly excluded by the AI pipeline, which recognises them as reconstruction noise rather than biological material. Intensity-based thresholding cannot make this distinction. The negative C2C mean for Salamander A (Specimen 2) shows that the pipeline produces tighter bone boundaries than binary thresholding, because it excludes the matrix halo that intensity-based methods incorporate due to the gradual density transition at the fossil-matrix interface. The AI-generated meshes are also natively aligned with the raw CT coordinate space, whereas the Avizo-generated meshes required mirror flipping and isotropic unit normalisation before registration could proceed. The Tier 3 C2C metrics for *Marmorerpeton wakei* (Specimens 1 and 2) reflect differences in segmentation completeness rather than poor accuracy of the AI output. Both specimens are multi-element vertebral complexes in which the binary thresholded reference mesh captures scattered matrix fragments, ghost trails, and noise as false positive bone, inflating the distance statistics. Where genuine shared anatomy is present, alignment between the two methods is tight: *Marmorerpeton wakei* (Specimen 1) has the best PPR coarse alignment in the entire assemblage (0.081 mm, 2.42 voxels), with sub-micron ICP corrections, confirming that the core skeletal structures are well-superimposed.

The total one-time investment is approximately 44 hours on a single NVIDIA A100 GPU (37-38 hours for SimCLR pre-training, 6 hours 10 minutes for U-Net training), after which per-specimen mesh generation requires only 1-3 minutes at inference. Manual segmentation of a single CT dataset typically takes on the order of 100 person-hours, and semi-automated workflows using commercial software still require persistent expert involvement for each specimen. Once trained, the framework can segment new CT datasets from Kilmaluag without retraining, preserving and accumulating training effort rather than discarding it after each session.

The external validation covers a meaningful range of phylogenetic and taphonomic diversity: stem salamanders (Salamander A: Specimens 1 and 2, *Marmorerpeton wakei*: Specimens 1 and 2), an early mammal (Mammaliaformes indet.), and an undescribed docodont, spanning distinct bone morphologies, body sizes, and preservation states. The pipeline handled the main taphonomic challenges of the Kilmaluag Formation successfully, including dolomitic matrix with near-identical radiodensity to bone, beam hardening, ring artefacts, and salt-and-pepper noise. Transfer to other formations, depositional environments, and scanning protocols is a natural extension for future work, as is systematic evaluation on invertebrate fossils, which present different preservation challenges including calcareous shells in limestone and silicified material in chert. Even in the scenario where retraining is required for a new depositional/lithological context, the framework’s zero-annotation requirement means the process is orders of magnitude less labour-intensive than conventional approaches: the approximately 44-hour GPU compute cost replaces hundreds of person-hours of manual annotation. Cross-domain generalization without retraining remains an open challenge in medical image segmentation, where acquisition protocols are far more standardised than in palaeontology; the best unsupervised domain adaptation methods close only 62% of the inter-domain performance gap on average.^48^. Progressive application of this pipeline to additional lithologies and scanning protocols will build toward broader generalization in future work.

Several methodological considerations should be noted. Binary thresholding in Avizo was chosen as the comparison baseline because it is the most widely used segmentation approach in palaeontological research and the only method that can be applied consistently across specimens without per-specimen manual intervention; even the AI-assisted modules in commercial platforms require manual annotation for each new specimen. The geometric comparison therefore evaluates how well a fully automated pipeline generalises across diverse specimens relative to the standard automated baseline it aims to replace. Three specimens were registered with ICP overlap set to 80% rather than 100% to prevent artefactual elements present only in the thresholded reference mesh, i.e., salt-and-pepper noise, matrix ghosting, and ring artefacts, from biasing the rigid alignment of core anatomy (see Methods). C2C signed distances for all six specimens were computed on the full, untrimmed point clouds regardless of the ICP overlap setting.

The current framework operates as a 2D slice-by-slice segmentation and does not enforce volumetric consistency between adjacent slices. In principle, 2.5D or native 3D architectures could improve inter-slice coherence, particularly for thin or obliquely oriented structures. In practice, however, training such architectures on palaeontological CT data is currently not feasible: volumetric approaches require high-quality 3D-annotated ground truth produced by expert manual segmentation of entire sub-volumes, and impose computational demands (GPU memory, training time, power consumption) that scale cubically with input size. Annotated 3D training data for fossils is essentially non-existent, and institutional computing resources are limited. The 2D approach adopted here is a pragmatic choice that balances segmentation quality against the realities of available data and computational access.

The pipeline has been trained and validated exclusively on vertebrate fossils; performance on invertebrate material, trace fossils, or heavily mineralised specimens is unknown and should be tested in future work. Slice-level classification metrics (Dice, IoU) were not computed for the six external validation specimens; geometric validation was performed at the mesh level only against manually thresholded references in CloudCompare v2 ^49,50^. We note that mesh-level agreement is the more relevant metric for downstream palaeontological applications (morphometrics, FEA, MDA), which operate on 3D surfaces rather than individual slices. The tight clustering of mesh-output Dice across taxonomically diverse external specimens (0.838–0.843 for five of six specimens, with Mammaliaformes indet. at 0.743 accounted for by ring artefacts in the source CT data) indicates limited variance of the taxonomically heterogeneous evaluation set, in contrast to the wider per-specimen spread reported for smaller fossil CT datasets evaluated across multiple backbone encoders^6^, suggesting that encoder choice is unlikely to be the dominant source of variation in this data regime.

This framework is designed to augment, not replace, palaeontological expertise. Conventional segmentation workflows consume the majority of expert time in the mechanical act of painting bone boundaries slice by slice — a task that contributes little to anatomical understanding and is prone to fatigue-induced error. By automating this phase, the framework frees the palaeontologist to focus where expertise is irreplaceable: evaluating biological plausibility, resolving anatomically ambiguous regions, and directing downstream quantitative analyses. The expert shifts from operator to analyst. Responsible deployment requires that automated outputs are treated as a starting point for expert review rather than a final answer, and this pipeline supports that: the model architecture, evaluation scripts, and masking pipeline are openly deposited, the training corpus and its taxonomic composition are fully reported, and all outputs are 3D meshes that can be visually inspected and corrected where required. Replacing black-box commercial platforms with an open, Python-based pipeline also improves transparency and lowers access barriers for researchers at under-resourced institutions.

By reducing per-specimen processing from approximately 100 person-hours to 6 hours +1-3 minutes, this pipeline makes large scale processing of entire CT collections tractable for the first time. Deployment of this pipeline at scale enables CT acquisition to feed directly into automated segmentation, producing analysis-ready 3D meshes deposited in open repositories such as MorphoSource^51^ and transforming existing CT data collections from static archives into dynamic, queryable research infrastructure. We release the framework as a resource for the community and anticipate that the dual-stream architecture described here, contrastive self-supervision combined with deterministic pseudo-labels will readily generalise to other scientific domains characterised by low-contrast imagery and scarce annotations.

## Methods

### Dataset assembly, preprocessing, and partitioning

A comprehensive dataset comprising 50,626 CT images of fossils, a resource assembled over 15 years of fieldwork at the Middle Jurassic Kilmaluag Formation, Scotland, forms the foundation for developing our segmentation framework (**Fig. 1**). The dataset represents a taxonomically diverse vertebrate assemblage, including fossil amphibians, archosaurs, pterosaurs, dinosaurs, and early mammals. These specimens are currently housed at National Museums Scotland (NMS) and CT volumes archived on MorphoSource (available on request). Scripts for the entire architecture were written using Python 3.11 and executed on BlueBear HPC (University of Birmingham high-performance computing cluster). The exact HPC hardware and runtime specifications are reported in **Supplementary Tables S3, S4 and S7**.

To ensure seamless integration with our framework, the core dataset was pre-processed by resizing all scans to a uniform resolution (224×224 px). Pixel intensity values are normalised to the range [0,1] and subsequently z-standardised (μ = 0.5, σ = 0.5) to maintain consistency across samples. The pre-processed dataset was subsequently partitioned using fixed seed pseudo-random sampling into training (19 datasets; 39,037 images) for SimCLR contrastive pretraining, validation (2 datasets; 3,765 images) for SimCLR checkpoint selection, and a U-Net partition (3 datasets, 7,824 images) for downstream segmentation training and evaluation. Within the U-Net partition, two specimens were used for training with rule-based deterministic masks, and one specimen of the same genus but a different individual was held out entirely as the internal test set; the Dice and IoU values reported in Results derive exclusively from this held-out specimen. The split CT image stacks, originally consisting of thousands of individual 2D TIFF files, were converted into Hierarchical Data Format (HDF5, .h5) to streamline data management and mitigate I/O bottlenecks. By consolidating these high-resolution TIFF slices into individual, self-describing containers, we eliminated the significant file-system overhead and directory-latency inherent in processing large image sequences. This conversion transformed the 2D data into a structured 3D volume, leveraging HDF5’s support for data chunking and ‘out-of-core’ processing. This conversion allowed our framework to perform high-speed random access on specific volumetric regions without exceeding RAM limits, while built-in lossless compression ensured a storage-efficient pipeline for downstream analysis.

### SimCLR pre-training on unlabelled fossil CT data

Palaeontological CT presents specific challenges for deep learning: sparsely labelled data, low signal-to-noise ratio, and radiographic artefacts including beam hardening, ring artefacts, and salt-and-pepper noise. Beyond these, the central technical obstacle is taphonomic density convergence, in which the mineral replacement of biological tissues during fossilisation produces X-ray attenuation coefficients in bone that are nearly indistinguishable from those of the surrounding rock matrix. This convergence is not noise that can be filtered out; it is a structural property of the data that pixel-intensity features alone cannot resolve. The SimCLR v1 framework was adopted specifically to address this (**Fig. 2**): by training a ResNet-50 encoder to extract textural and structural signatures rather than intensity statistics, the model learns to discriminate fossil from matrix in exactly the regime where thresholding fails. Contrastive pre-training is performed on a large corpus of unlabelled images before fine tuning for pixel-wise classification.

To capture high inter subject variability (i.e., intrinsic physical and radiographic differences among fossil specimens resulting from microscale taphonomy, morphological diversity, scan quality and artefacts, structural deformation and fragmentation), and complex preservation states, we implemented a domain specific stochastic data augmentation pipeline. For each CT slice, the framework generates a positive pair of augmented images (*x_i_*and *x_j_*), through random geometric and intensity-based operations. These include random resizing /cropping (scale = 0.6–1.0), vertical flipping (p=0.5), horizontal flipping (p=0.2), random rotation (±15°, p = 0.3), Gaussian blur (p=0.3) to learn global anatomical organisation and local bone micro-texture, as well as arbitrary fossil orientation within the matrix. Intensity and contrast jitter (brightness: p=0.8, contrast: p=0.8) enforce invariance to physical constraints of X-ray imaging. Pre-training acts as a regulariser, mapping noisy and clean inputs to proximate latent representations, thereby improving robustness to stochastic graininess and structural distortions typical of high resolution µCT data. Leveraging taxonomically diverse images in our partitioned dataset, this approach yields domain specific representations better suited to palaeontological segmentation than large-volume generalised image datasets (e.g., ImageNet). The feature representations (*h_i_* and *h_j_*) extracted by the ResNet50 base encoder are passed through a ‘projection head’ to map them into a latent space where the contrastive loss was applied. This projection head consists of a multi-layer perceptron (MLP) featuring two linear layers separated by a Rectified Linear Unit (ReLU) non-linear activation function. Mapping the representations to this latent space was critical as it prevents the loss of important structural information that might occur if the contrastive objective were applied directly to the feature vectors used for downstream segmentation. The model training was assessed using the NT-Xent loss, which encourages the network to maximize the agreement between the positive pair in the latent space while simultaneously pushing them away from all other images in the training batch. To monitor the effectiveness of the self-supervised pre-training, we tracked both the average NT-Xent loss and Contrastive Accuracy across 250 training epochs (batch size= 64, learning rate = 3×10⁻^4^, weight decay: 1 × 10⁻⁴).

Validation of the learned representations was conducted through two primary ‘sanity check’ visualisations, Grad-CAM and UMAP, to ensure the model was capturing meaningful biological structures rather than background noise. While Grad-CAM produces heatmaps that highlight the regions of the CT slice most influential to the model’s feature extraction, and UMAP applies non-linear dimensionality reduction to the high-dimensional feature vectors to visualise the structure of the dataset in 2D Cartesian space.

### Rule-based pipeline for reproducible spatial ground truth

A fully deterministic, rule-based pipeline generated coarse binary masks of fossilised bone from each CT slice, providing standardised spatial guidance for subsequent U-Net training. The pipeline operates through five sequential stages: (1) intensity normalisation via percentile clipping to suppress outliers and enhance the radiodensity range of mineralised bone; (2) Gaussian and median blurring to mitigate high-frequency noise and radiographic artefacts; (3) seed identification using Otsu thresholding^52^ followed by (4) outward region expansion to capture the full spatial extent of bone within each slice; morphological refinement through top-hat transformations and erosion/dilation cycles to fill internal gaps, remove disconnected artefactual fragments, and sharpen edges; and (5) connected component filtering to retain only regions exceeding a minimum physical area threshold and overlapping with identified bone seeds. All parameters were determined empirically on the training partition and held fixed thereafter. Given identical input, the algorithm produces identical output. Full implementation is available in the code repository.

### Dual-stream knowledge transfer to U-Net

In the final segmentation phase, we employed a knowledge transfer strategy^53^ that integrated domain-specific feature representations derived from self-supervised SimCLR v1 pre-training with the standardised guidance of automatically generated ground-truth masks. By initialising the U-Net with weights from the SimCLR ResNet-50 encoder, the model began with an advanced representation of fossilised bone textures and structural signatures that differentiated them from surrounding geological noise. These learned feature representations, combined with rule-based automated ground-truth masks, provided consistent and deterministic delineations of bone specimens. This dual transfer learning stream ensures that the U-Net did not rely exclusively on raw pixel intensities, which are often unreliable due to taphonomic density convergence, but instead integrated high-level textural cues with the standardised spatial constraints defined by the automated masking algorithm.

The core segmentation architecture consisted of a modified U-Net optimised for high-resolution volumetric data. In contrast to a conventional U-Net, this configuration incorporated a deeper encoder pathway (e1–e4) with progressively increasing channel depths (64-2048), enabling the extraction of increasingly abstract morphological features (Fig. 1). The central bottleneck was substantially expanded, incorporating a 7×7 convolutional block with integrated ReLU activation functions to model complex spatial relationships at a 7×7 resolution. The decoder pathway (up4–up1) symmetrically mirrored the encoder to reconstruct the segmentation map, while skip connections preserved fine-scale structural detail, including delicate fossil boundaries. During training across 500 epochs, the model employed a composite Dice and Cross-Entropy loss function to address the pronounced class imbalance between bone and matrix. Further optimization was carried out using AdamW^54^, which is widely considered highly efficient for deep neural networks, while the OneCycleLR^51^scheduler was applied to rapidly identify an optimal learning rate during training. Training convergence metrics are shown in **Fig. 3a, b**. Model performance was finally evaluated on the held-out internal test specimen, which was not used during U-Net training.

### 3D mesh generation and geometric comparison

The final stage of the pipeline involved reconstructing volumetric 3D meshes from the 2D segmentation masks generated by our modified U-Net model. Six external fossil CT datasets, encompassing stem salamanders (Salamander A.: Specimens 1 and 2; MorphoSource: 00084381 and high-resolution rescan of 000071513 respectively), *Marmorerpeton wakei* (Specimens 1 and 2; MorphoSource: 000700518 and 000700519), early mammals: (Mammaliaformes indet.; MorphoSource: 000693738), and an undescribed docodont (MorphoSource: 000721884), were segmented.

This process began by stacking the 2D masks to form a 3D volume that preserved the shape and spacing of the original CT scans. To ensure structural integrity before surface extraction, the 3D volume underwent a multi-step refinement process comprising region growing from high-confidence bone voxels, intensity filtering to exclude non-bone regions based on CT grayscale values, and morphological operations to connect bone fragments and eliminate noise.

Following these refinement steps, the marching cubes algorithm^55^ was employed to convert the 3D volume into a triangulated surface mesh using an iso-level of 0.5. This resulted in watertight 3D meshes suitable for further analysis, visualization, or 3D printing. The quality of the output meshes was assessed through quantitative metrics, including the count of STL vertices and faces, as well as mean Dice and IoU scores derived from the underlying segmentation. Across the evaluated specimens, the mesh generation demonstrated run times generally taking between 1-3 minutes per specimen.

Additionally, the same six fossil CT datasets were segmented in ThermoScientific Avizo (ver. 2024.2) using binary thresholding. The resulting meshes were compared with those generated by our end-to-end framework through three quantitative evaluation steps in CloudCompare v2 (**Fig. 4**). First, point-pair registration (PPR)^43^ followed by a coarse Iterative Closest Point (ICP)^44^ alignment was applied to correct arbitrary initial orientations and bring the meshes into approximate correspondence. The target was a root mean-squared (RMS) error within a few multiples of the voxel size, ensuring sufficient proximity for fine-scale optimisation. Second, a scale-calibrated ICP^44^ was performed to further optimise alignment by iteratively minimising nearest-neighbour distances while allowing for minor global scale differences. A satisfactory fit was defined by a scale factor close to 1.0, indicating high dimensional fidelity, and a mean cloud-to-cloud (C2C)^45^ distance approaching zero, reflecting near-identical overall surface geometry. Finally, C2C distance and Hausdorff analyses were conducted to quantify local deviations and extreme discrepancies. A low standard deviation indicated consistently tight surface agreement, whereas a high one-sided Hausdorff distance highlighted localised mismatches (such as small bones or noise artefacts) present in one mesh but absent in the other. Together, this three-stage evaluation framework ensured not only robust global alignment but also rigorous statistical assessment of biologically meaningful features, enabling clear discrimination between the genuine anatomical signal and manual segmentation artefacts.

For Salamander A (Specimen 1), Salamander A (Specimen 2) and Docodonta (undescribed), ICP overlap was set to 100%, meaning all points were used and the resulting RMS captures full geometric divergence. For Mammaliaformes indet., *Marmorerpeton wakei* (Specimens 1 and 2), overlap was set to 80%, computing correspondences on only the best-matching 80% of points and discarding the worst 20% as outliers during each iteration. This trimmed setting was applied to specimens exhibiting significant non-overlapping geometry: multi-element vertebral complexes and fragmented specimens with prominent ring artefacts, where outlier fragments present exclusively in the thresholded reference mesh would otherwise bias the rigid alignment of core anatomy. The C2C signed distances in the third phase were computed on the full, untrimmed point clouds for all six specimens regardless of the ICP overlap setting.

## Data availability

The CT datasets analysed in this study are derived from fossil specimens housed at National Museums Scotland (NMS). Raw CT volumes are accessioned and deposited in MorphoSource^51^ (www.morphosource.org), subject to NMS institutional access procedures. Owing to file size, the processed dataset (HDF5 image stacks and segmentation masks used for SimCLR pre-training and U-Net training) is archived in the University of Birmingham Research Data Store (RDS). 4K rendered animations of the segmented test specimens are archived on Zenodo [DOI: 10.5281/zenodo.20579028]. Segmented meshes (.STL/.OBJ files) and the processed-data archive are available from the authors on reasonable request.

## Code availability

The repository hosted at https://github.com/roy-arindam-1991/Simclr-v1-50k-automated-pipeline; under a GNU General Public License v3.0 and archived on Zenodo [DOI:10.5281/zenodo.20242346] contains: the U-Net and SimCLR model architecture definitions; the training loop and evaluation scripts; the rule-based deterministic masking pipeline; the 3D mesh generation and CloudCompare registration workflows; and a full environment specification (Python version, package dependencies, CUDA version). Hyperparameter values optimised during training and pre-trained model weights are not included in the public repository; these are available from the authors on request, subject to a brief description of intended use.

## Acknowledgements

The authors thank National Museums Scotland for access to the fossil specimens. This work used the BlueBEAR High Performance Computing (HPC) facility at the University of Birmingham, and we thank Kiran Pandurang Phalke and Thomas Kappas (Research Software Group, Advanced Research Computing, IT Services, University of Birmingham) for training on HPC usage and code unit testing.

## Author Contributions

A.R. and R.J.B. conceived and designed the study, which formed the basis of a Horizon Europe Marie Skłodowska-Curie Actions Postdoctoral Fellowship proposal. A.R. led the methodology and analysis, and wrote the original draft. P.G. contributed to the pipeline architecture, training code, and data preprocessing. F.W. assisted with cleaning and processing the meshes in Blender. B.H., A.S.-J. and S.T.S.P. supervised A.R. during the AI secondment at the Natural History Museum, London, and provided feedback on pipeline development. S.C.R.M. organised the AI secondment to the Natural History Museum, London, and advised on the structure of the manuscript. R.J.B. supervised the project and provided specimen access. All authors reviewed, edited and approved the final manuscript.

