## Supplementary Information for "Breaking the bottleneck: self-supervised deep learning framework for fully automated fossil CT segmentation"

**Contents:**

*Supplementary Figures* (**Fig. S1-S3**)

*Supplementary Notes* (**1-6**) *& Tables* (**S1-S15**)

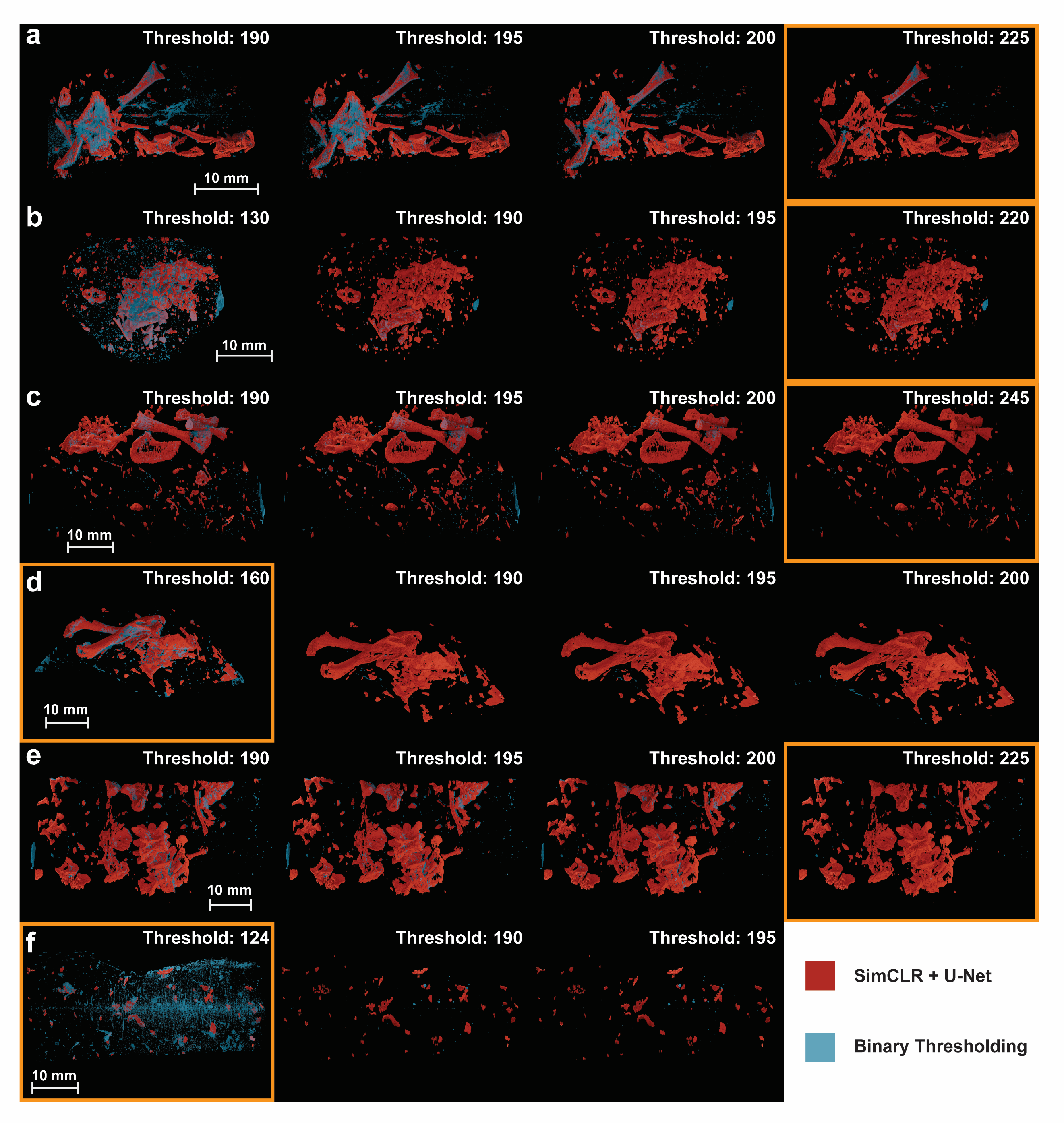

***Fig. S1 Threshold sensitivity analysis for binary segmentation across all six external validation specimens.*** *Each row shows successive binary threshold levels applied to the CT volume of the labelled specimen, with the SimCLR + U-Net prediction (red) overlaid for comparison against the manual binary thresholding output (blue). Threshold values increase from left to right within each row. The optimal binary threshold — defined as the highest value at which skeletal elements remain coherent with minimal noise and before cortical surfaces begin to dissolve, is indicated by a gold box.* ***(a)*** *‘Salamander A’ Specimen 1 (MorphoSource 00084381; voxel size 0.0154 mm); optimal threshold: 225.* ***(b)*** *‘Salamander A’ Specimen 2 (rescan of MorphoSource 000071513; voxel size 0.0228 mm); optimal threshold: 220.* ***(c)*** *Marmorerpeton wakei Specimen 1 (MorphoSource 000700518; voxel size 0.0336 mm); optimal threshold: 245.* ***(d)*** *Marmorerpeton wakei Specimen 2 (MorphoSource 000700519; voxel size 0.0325 mm); optimal threshold: 160.* ***(e)*** *Docodonta indet. (MorphoSource 000721884; voxel size 0.0252 mm); optimal threshold: 225.* ***(f)*** Mammaliaformes *indet. (MorphoSource 000693738; voxel size 0.0243 mm); optimal threshold: 124. For (f) Avizo mesh export (STL) failed beyond threshold 195 owing to insufficient points remaining in the segmented bone volume. Scale bars: 10 mm.*

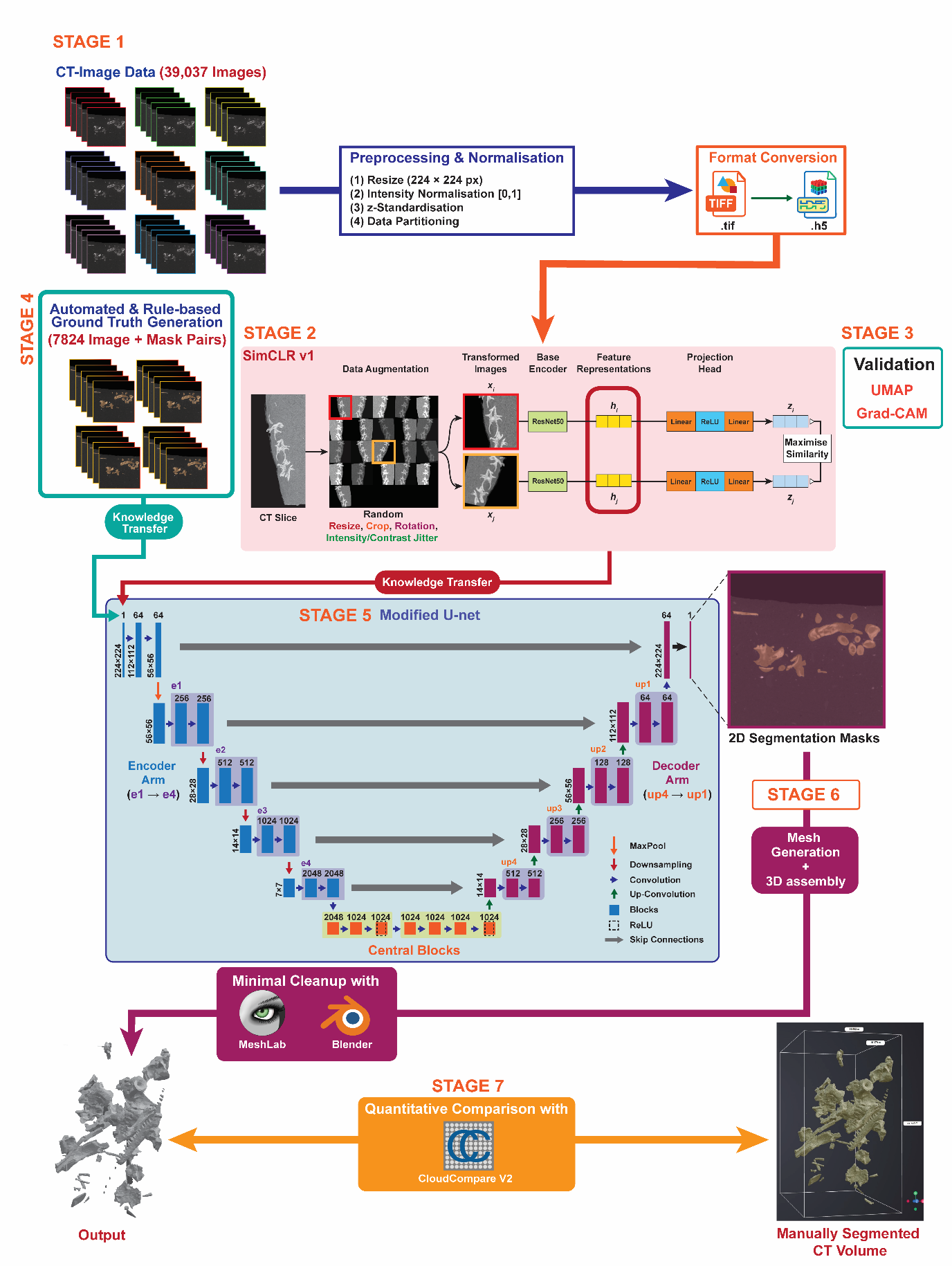

***Fig. S2 Schematic of the seven-stage SimCLR + U-Net pipeline for automated fossil CT segmentation.*** *Each stage is colour-coded to its corresponding workflow panel and cross-referenced to its Supplementary Note. Stage 1 (****Supplementary Note 2****; blue panels): raw CT slices undergo resizing, normalisation, z-standardisation, specimen-level partitioning, and consolidation from TIFF into HDF5. Stage 2 (****Supplementary Note 3****; pink panel): SimCLR v1 self-supervised pretraining, in which augmented positive pairs (xᵢ, xⱼ) are passed through a shared ResNet-50 encoder to generate feature representations (hᵢ, hⱼ) and projection-head outputs (zᵢ, zⱼ) optimised under the NT-Xent contrastive loss. Stage 3 (****Supplementary Note 3****; representative outputs in* ***Fig. S3*** *and* ***Fig. 2c*** *of the main text): encoder feature quality is validated using UMAP clustering and Grad-CAM heatmaps. Stage 4 (****Supplementary Note 3****; green panel): automated rule-based ground-truth masks (7,824 slices) are generated through thresholding, Otsu seed detection, region growing, and morphological refinement, eliminating manual annotation. Stage 5 (****Supplementary Note 4****; mint panel): the SimCLR-pretrained encoder weights are transferred (red arrow) into a modified U-Net, with skip connections (grey arrows) bridging encoder and decoder to retain fine-grained boundary information and produce predicted 2D segmentation masks. Stage 6 (Supplementary Note 5; purple panel): predicted masks are stacked into a 3D volume, from which the marching cubes algorithm extracts a triangulated surface mesh, with minimal cleanup in MeshLab and Blender; per-specimen mesh outputs are reported in Table 1 of the main text. Stage 7 (Methods of the main text; orange panel): AI-generated meshes are compared to manually segmented reference meshes via a three-phase registration protocol in CloudCompare v2, with validation statistics reported in* ***Table 2*** *of the main text. The green Knowledge Transfer arrow indicates the flow of image–mask training data from Stage 4 to Stage 5; the red Knowledge Transfer arrow indicates the transfer of SimCLR-pretrained encoder weights from Stage 2 to Stage 5.*

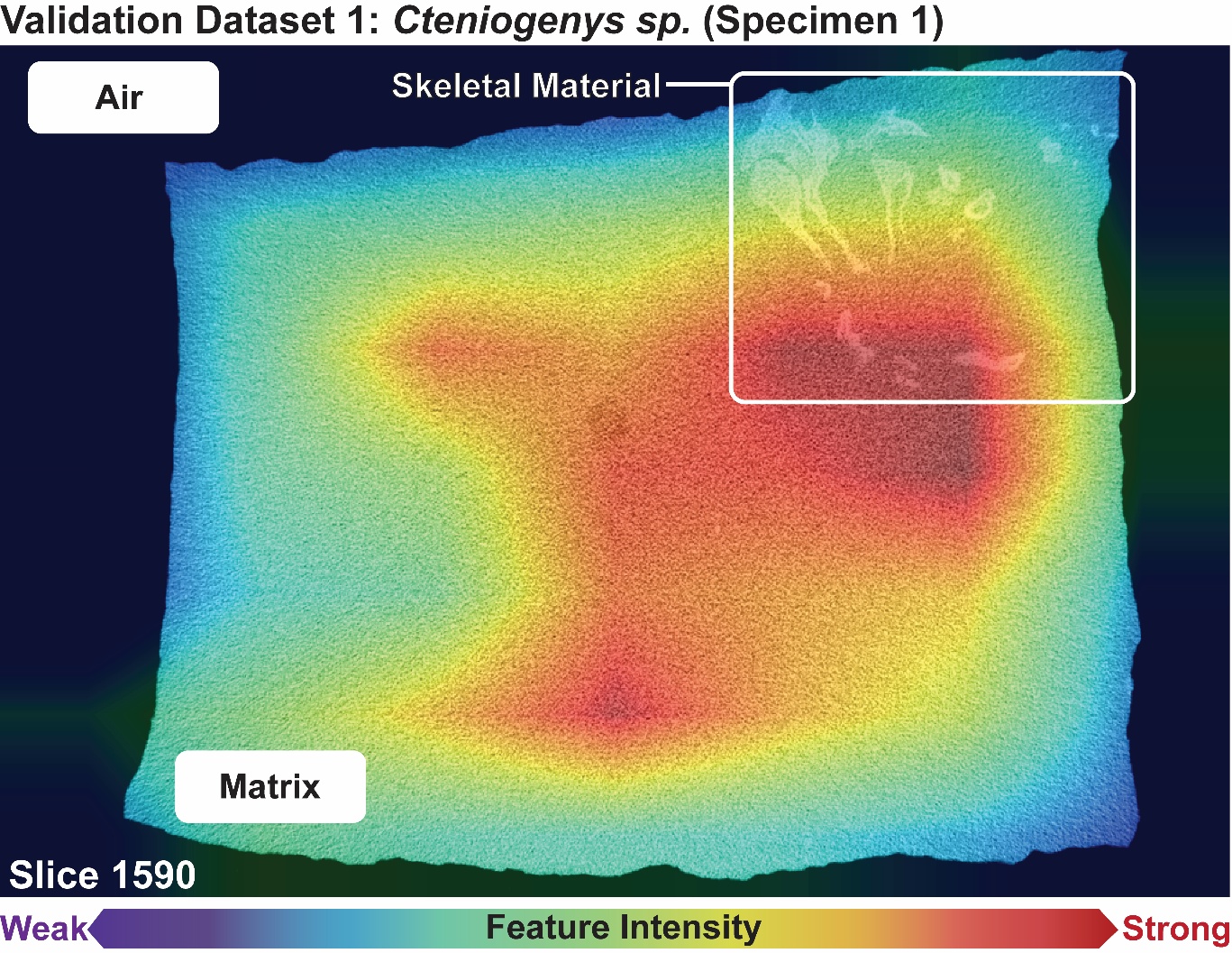

***Fig. S3 Grad-CAM feature activation map for a representative CT slice of Cteniogenys sp. (Validation Dataset, Specimen 1).*** *Gradient-weighted Class Activation Mapping (Grad-CAM) heatmap overlaid on slice 1590, illustrating the spatial distribution of feature intensity learned by the SimCLR-pretrained ResNet-50 encoder. Colour scale ranges from weak activation (purple–blue) to strong activation (yellow–red). Three material phases are annotated: air (peripheral dark regions, minimal activation), sedimentary matrix (mid-field, intermediate activation), and skeletal material (upper right, enclosed by white box, peak activation). The strongest activations (red) are concentrated within and immediately surrounding the skeletal elements (outlines are deliberately drawn as darker contours within the high-intensity zone for ease of comprehension), confirming that the encoder has learned to prioritise bone-like attenuation signatures over matrix and air. The broad intermediate-activation zone across the matrix reflects the taphonomic density convergence characteristic of Kilmaluag Formation specimens, in which diagenetic mineralisation produces attenuation values that partially overlap with bone-the principal segmentation challenge addressed by this framework. The spatial correspondence between peak Grad-CAM activation and confirmed skeletal material provides independent, slice-level validation that the network's learned representations are anatomically meaningful rather than artefactual.*

**Supplementary Note 1**

***Supplementary Table S1*** *A detailed comparison of features among select segmentation software and the AI architecture presented in this study*

| **Feature** | **Software** | | | | | | | | | | | | | |
| --- | --- | --- | --- | --- | --- | --- | --- | --- | --- | --- | --- | --- | --- | --- |
|  | **Dragonfly** | **Avizo** | | **Mimics** | | **VGStudio** | | **Biomedisa** | | | **SPIERS** | | **3D Slicer** | **Roy et al. (this study)** |
| **Developer** | [Comet Technologies Canada (Object Research Systems)](https://dragonfly.comet.tech/en/resources/user-manuals) | [Thermo Fisher Scientific](https://documents.thermofisher.com/TFS-Assets/MSD/Software-Updates/amira-avizo3d-20242-release-notes.pdf) | | [Materialise NV](https://www.materialise.com/en/healthcare/mimics/ai-enabled-segmentation) | | [Hexagon (Volume Graphics GmbH)](https://volumegraphics.hexagon.com/en/products/vg-software-release-updates.html) | | ANU CTLab^1^ | | | Sutton et al.^2^; Palaeoware community | | Kitware, BWH, community | Roy et al., University of Birmingham |
| **Licence / Cost** | [FreeD licence](https://dragonfly.comet.tech/en/products/license-models/freed-license) (restricted: excludes gov’t, non-profit, [5 territories,](https://dragonfly.comet.tech/en/products/license-models/freed-license/freed-country-restrictions) HPC/cloud). Commercial licence required otherwise | Commercial only. No free academic programme | | Commercial only. Research version available in some regions | | Commercial only. VGTRAINER module at additional surcharge | | Free, open-source (Apache 2.0) | | | Free, open-source (GPL v3) | | Free, open-source (BSD-style) | Free, open-source (GPL v3) |
| **Primary design domain** | Materials science, industrial CT, general biomedical | Materials science, life sciences, geosciences | | Clinical medical imaging (orthopaedic, maxillofacial) | | Industrial CT, non-destructive testing, manufacturing QA | | Biomedical (insects, microanatomy, some fossil) | | | Serial grinding tomography (invertebrate palaeontology) | | Clinical medical imaging (human anatomy)^3^ | Vertebrate fossil micro-CT with complex taphonomic signatures |
| **Latest version / update cadence** | 3D World 2025. ~Annual releases | v2025.1 (Apr 2025). ~2 releases/year | | v28.0 (2025). ~Annual releases | | v2026.1; VGTRAINER added Apr 2025. ~Annual | | Active on GitHub (2025). Irregular, academic-driven | | | v3.1.0 (Aug 2024). Sparse: multi-year gaps between releases. No AI roadmap | | v5.10 (2025). ~Quarterly. AI extensions depend on external projects | v1.0 (2026). Under active development with GPL v3 codebase on GitHub |
| **AI/ML segmentation capability** | Segmentation Wizard: Random Forest on handcrafted texture features + basic U-Net.^4^ Supervised, per-specimen | Segmentation+ Workroom ([v2024.2](https://documents.thermofisher.com/TFS-Assets/MSD/Software-Updates/amira-avizo3d-20242-release-notes.pdf)): [U-Net with VGG16, VGG19, or ResNet-18](https://www.thermofisher.com/software-em-3d-vis/customerportal/news/amira-avizo-2024-2-advanced-ai-features-and-exciting-enhancements/) backbones. Supervised, per-specimen | | 9 pre-trained ‘[AI-enabled’ algorithms for human bony anatomy (hip, torso, skull](https://www.materialise.com/en/healthcare/mimics/ai-enabled-segmentation)). Locked models, not adaptive. No user-trainable DL | | [VGTRAINER (v2025.1)](https://volumegraphics.hexagon.com/en/products/vgtrainer.html): user-trained DL models for industrial segmentation. Requires manually labelled training data | | [Smart Interpolation](https://github.com/biomedisa/biomedisa/blob/master/README/smart_interpolation.md) (random walks from sparse manual labels) + supervised 3D U-Net requiring fully labelled volumes | | | None. [Threshold-based and manual mask painting only](https://github.com/palaeoware/SPIERS) | | Extensions: [TotalSegmentator](https://discourse.slicer.org/t/monai-and-totalsegmentator/36636) (104 human structures), MONAI Auto3DSeg, MONAILabel. All pre-trained on human clinical data or require user-labelled training data^5^ | SimCLR v1 self-supervised contrastive pre-training (ResNet-50 backbone) + deterministic pseudo-label generation + modified U-Net.  Zero manual annotation |
| **Manual annotation required?** | Yes. Per-specimen manual ROI painting for every new dataset | Yes. 2–5 manually annotated ROIs per specimen | | No (pre-trained), but models are fixed to human anatomy; not trainable on fossils | | Yes. Manually labelled training datasets required | | Yes. Sparse pre-segmented slices (interpolation) or fully labelled volumes (DL) | | | Yes. Fully manual, slice-by-slice | | TotalSegmentator: no (but human-only). Custom training: yes, requires labelled data | No. Entire pipeline is annotation-free. Pseudo-labels generated deterministically |
| **Self-supervised / contrastive pre-training** | No | No | | No | | No | | No | | | No | | No | Yes. SimCLR v1 on 39,037 unlabelled fossil CT slices. Learns domain-specific representations (taphonomic noise, beam hardening, density gradients) |
| **DL backbone architecture** | Random Forest + basic U-Net (shallow encoder) | VGG16, VGG19, or ResNet-18 (max 18 layers) | | Undisclosed proprietary models | | Undisclosed (user-trained via VGTRAINER) | | Standard U-Net (no pre-trained encoder) | | | N/A (no DL) | | Varies by extension; typically, nnU-Net or ResNet-based | ResNet-50 (50 layers; substantially deeper and more expressive) pre-trained via SimCLR, feeding modified U-Net with deeper encoder path |
| **Training signal preserved across specimens?** | No. Classifier is specimen-specific and discarded after each session | | No. Per-specimen training discarded | | N/A (fixed pre-trained models) | | Yes, within a trained model for identical part types | | DL model can be reused on similar specimens; but requires full annotation to create | No. Each specimen starts from zero | | Pre-trained models reusable within their domain (human anatomy) | | Yes. SimCLR encoder accumulates representations across 50,626 taxonomically diverse images. Training effort compounds, not discards |
| **Batch processing / HPC scalability** | FreeD: HPC/cloud prohibited. Commercial: local workstation GUI only | | GUI-based, single workstation. Some Python scripting possible | | Cloud segmentation available (Mimics Flow). Desktop: GUI-only | | VGinLINE for production; primarily GUI-based | | Web-based or local. No native HPC batch pipeline | GUI-only. No scripting, no batch capability | | Python-scriptable. Extensions vary in HPC support | | Command-line, fully scriptable, parallelisable. Deployable on HPC clusters, cloud, and containerised pipelines. 1–3 min/specimen at inference |
| **Fossil CT-specific features** | None. General-purpose | | None. General-purpose | | None. Human anatomy only | | None. Industrial manufacturing | | Used for Little Foot skull^6^) and amber^1^ inclusions, but via manual pre-segmentation, not autonomous DL | Originally designed for serial grinding of Herefordshire invertebrates^2^; adapted for CT but no fossil-specific automation | | None. Pre-trained on human clinical CT | | Trained on 50,626 fossil micro-CT images spanning amphibians, reptiles, dinosaurs, early mammals. Handles beam hardening, isodense bone/matrix, taphonomic noise, incomplete preservation |
| **Transparency/ reproducibility** | Proprietary. Algorithmic details, feature selection, parameterisation obscured | | Proprietary. Black-box models | | Proprietary. ‘Locked and validated before release’; no user access to model internals | | Proprietary. VGTRAINER trains user models but core code is closed | | Open-source. Random walk algorithm published | Open-source (GPL v3). Fully transparent | | Open-source (BSD). Extensions vary | | Open-source (GPL v3). Deterministic: same input always produces identical output. All parameters, architecture, training corpus reported. Code on GitHub, weights on request |
| **Published fossil CT performance** | Edie, et al. ^7^ accuracy 0.97 but Dice only 0.13 on bivalves using Dragonfly Segmentation Wizard (5 training slices). Low Dice reflects poor overlap despite high pixel accuracy due to class imbalance | | Nešić, et al. ^8^: IoU 0.462 (electron microscopy, not fossil CT) | | None on fossil material | | None on fossil material | | Little Foot skull segmentation (semi-automated interpolation, not DL) | No performance benchmarks published | | None on fossil material | | Dice 93.66%, IoU 82.42% (held-out specimen). Sub-voxel mesh agreement on 6 external specimens across 4 higher-level taxa |
| **Key limitation for fossil CT** | FreeD excludes museums, govt. labs, 5 territories. ML is supervised, shallow, specimen-specific | | Expensive. Supervised DL with shallow backbones. No fossil-specific training | | Locked to human anatomy. Cannot be retrained on fossil data. Expensive | | Designed for industrial QA (porosity, inclusions). No palaeontological application. Expensive | | Requires manual annotation (sparse or full). Standard 3D U-Net without contrastive pre-training. No fossil-specific representations | Entirely manual. No AI/ML. Slow for large vertebrate CT volumes. Originally designed for invertebrate serial grinding | | Pre-trained models locked to human clinical anatomy. Custom training requires labelled fossil data (which is the bottleneck this paper addresses) | | Validated on single formation (Kilmaluag). Cross-formation transfer identified as future work. Segments bone from matrix as unified object (element separation is downstream) |

**Supplementary Note 2**

***Data preprocessing and dataset partitioning.***

CT datasets comprised single-layer TIFF-format slices of fossilised bone specimens. All images were resized to 224 × 224 pixels and pixel intensities normalised to [0, 1], followed by standardisation to a mean of 0.5 and standard deviation of 0.5. This pipeline ensured consistent input representation and stable optimisation across specimens acquired on different scanners and at different voxel resolutions.

The full dataset of 24 MorphoSource deposited specimens (50,626 slices) was partitioned at the specimen level prior to any model training to prevent data leakage between sets. Subfolders were randomly shuffled using a fixed seed (42) to ensure reproducibility. Partitioning is summarised in ***Supplementary Table S2***. Specimens for SimCLR pretraining were selected from the Kilmaluag Formation holdings on MorphoSource on the basis of scan completeness, voxel resolution, and absence of severe beam hardening artefacts. The full training specimen list will be released alongside the model weights and training code upon publication. The six external validation specimens on which all geometric validation claims rest are fully disclosed in **Tables 1–2** of the main text, with MorphoSource accession numbers provided therein.

***Supplementary Table S2*** *Dataset partitioning summary.*

| **Task** | **Specimens (n)** | **Slices** | **Proportion** |
| --- | --- | --- | --- |
| SimCLR pretraining | 19 | 39,037 | 77.1% |
| SimCLR validation | 2 | 3,765 | 7.4% |
| U-Net training+ held out dataset testing | 3 (2+1) | 7,824 | 15.5% |
| **Total** | **24** | **50,626** | **100%** |

***Data storage, format conversion, and computational infrastructure.***

Following dataset partitioning (**Supplementary Note 2**), all CT image stacks and corresponding segmentation masks were converted from TIFF format into Hierarchical Data Format version 5 (HDF5; .h5)^9^, as illustrated in the Format Conversion step (**Fig. 1**, **S2**, Stage 1). The rationale for this format conversion is described in the Methods section.

Two distinct computational jobs correspond to this stage of the workflow. The mask generation job (mask_gen.sh) produced the 7,824 image–mask pairs used for U-Net training, corresponding to the Automated and Rule-based Ground Truth Generation step (**Fig. 1**, **S2**, Stage 4). The HDF5 conversion job (mask_h5.sh) subsequently consolidated those outputs into the structured HDF5 format shown in the Format Conversion step (**Fig. 1**, **S2**, Stage 1). All jobs were executed on the BlueBEAR high-performance computing cluster at the University of Birmingham. Computational resource usage for each step is summarised in Supplementary **Table S3**.

***Supplementary Table S3*** *Computational resource usage for data preparation steps, corresponding to the Preprocessing and Normalisation and Format Conversion stages of* ***Fig. 1*** *(Stage 1).*

| **Workflow Step** | **Description (as shown in Fig 1)** | **CPUs** | **Memory** | **Runtime** |
| --- | --- | --- | --- | --- |
| Dataset splitting | Preprocessing & Normalisation (step 4) | 1 | 16 GB | ~40 s |
| Full HDF5 pipeline | Format Conversion (TIFF → .h5) | 1 | 16 GB | ~41 min |
| Mask generation (mask_gen.sh) | Automated & Rule-based Ground Truth Generation | 18 | 128 GB | 3 min 19 s |
| HDF5 conversion (mask_h5.sh) | Format Conversion (TIFF → .h5) | 16 | 128 GB | 27 s |

All jobs were CPU-only; no GPU allocation was required for data preparation. Node-level CPU specifications follow standard BEAR cluster configuration (Intel Xeon architecture).

**Supplementary Note 3**

***SimCLR v1 contrastive pretraining.***

The HDF5-formatted dataset of 39,037 CT slices (***Supplementary Note 2***) was used to train a SimCLR v1 self-supervised model, corresponding to the SimCLR v1 stage (**Fig. 1**, **S2**, Stage 2). The ResNet-50 backbone was modified to accept single-channel greyscale CT images, with a projection head comprising a linear layer, ReLU activation, and a final linear layer projecting to a 128-dimensional latent space. The model was trained using the NT-Xent contrastive loss with a temperature^10^ of 0.2. To promote encoder invariance to the imaging variability inherent in fossil CT data, greyscale-specific augmentations were applied during training: random resized cropping (scale 0.6–1.0), horizontal flipping (p = 0.5), vertical flipping (p = 0.2), random rotation ±15° (p = 0.3), and Gaussian blur (p = 0.3). Mixed-precision training (fp16) was enabled on GPU^11^. Reproducibility was ensured with a fixed seed of 42. Model checkpoints were saved after every epoch, with the best-performing checkpoint retained for knowledge transfer to the U-Net training stage (Stage 5; Supplementary Note 4). Full training configuration, HPC specifications, and performance outcomes are summarised in ***Supplementary Table S4***.

***Supplementary Table S4*** *SimCLR v1 training configuration, HPC specifications, and performance metrics.*

| **Parameter** | **Value** |
| --- | --- |
| HPC cluster | BlueBEAR (University of Birmingham) |
| Job scheduler | SLURM |
| GPU | 1 × NVIDIA A100 |
| CPU cores | 18 |
| RAM | 120 GB |
| Maximum runtime allocated | 48 hours |
| Estimated actual runtime | ~37–38 hours |
| PyTorch version | 2.1.2 (CUDA 12.1) |
| Python version | 3.11 (GCC 12.3.0) |
| Key dependencies | torchvision 0.16.0, scikit-image 0.22.0, h5py 3.9.0 |
| Training slices | 39,037 |
| Epochs | 250 |
| Batch size | 64 (gradient accumulation ×16; effective batch 1,024) |
| DataLoader workers | 4 |
| Training accuracy | 0.9389 |
| Training loss | 0.2036 |
| Internal validation accuracy | 0.9366 |
| Internal validation loss | 0.2312 |

***Representation validation***

Following SimCLR pretraining, the quality of the learned feature representations was assessed prior to knowledge transfer, corresponding to the Representation Validation stage (***Fig. 1, S2***). UMAP dimensionality reduction^12^ was applied to the 2,048-dimensional feature vectors extracted by the trained ResNet-50 encoder to evaluate whether bone and matrix embeddings form spatially distinct clusters in feature space without supervision. Grad-CAM heatmaps^13^ were generated to provide spatially explicit, slice-level confirmation that peak activations localise to skeletal material rather than surrounding matrix or air. Together, these two analyses confirm that the encoder has learned anatomically meaningful representations before any labelled data are introduced. Representative outputs are shown in ***Fig. 2c*** of the main text and ***Fig. S3***.

***Rule-based ground-truth mask generation.***

Deterministic segmentation masks for the U-Net training corpus (7,824 image–mask pairs) were generated automatically using a rule-based algorithm, corresponding to the Automated and Rule-based Ground Truth Generation stage (***Fig. 1****,* ***S2****, Stage 4*) The algorithm combined bone density percentile thresholding (clipping at 0.1–99.9^th^ percentile to suppress outliers), Gaussian and median blurring for noise reduction, Otsu-based seed detection followed by fixed-threshold region growing, morphological refinement (top-hat transformation, erosion, dilation)^14^, and connected component filtering to retain only regions overlapping seed detections and exceeding a minimum physical area threshold. Voxel spacing was extracted per specimen from CSV metadata to convert physical thresholds to pixel-space equivalents. Key algorithm parameters are summarised in ***Supplementary Table S5***.

***Supplementary Table S5*** *Rule-based mask generation parameters*

| **Parameter** | **Value** | **Description** |
| --- | --- | --- |
| Physical area threshold | 0.15 mm² | Minimum bone area retained per connected component |
| Kernel size | 41 | Controls smoothing extent during noise reduction |
| Growth threshold | 45 | Controls outward mask expansion from seed regions |
| Percentile clip range | P_0.1_–P_99.9_ | Suppresses intensity outliers prior to thresholding |

**Supplementary Note 4**

***U-Net segmentation training***

The SimCLR-pretrained ResNet-50 encoder weights were transferred to a modified U-Net architecture^15^ via the Knowledge Transfer step (***Fig. 1, S2,*** *Stage 5*). The encoder arm (e1–e4) preserves the ResNet-50 feature hierarchy; skip connections bridge encoder and decoder to retain fine-grained boundary information; and the decoder arm (up4–up1) progressively reconstructs spatial resolution to the original 224 × 224 input. The U-Net architecture is described in ***Supplementary Table S6***. Training used a composite Dice and Weighted Cross-Entropy loss^16^, AdamW optimisation^17^, and an OneCycleLR learning rate scheduler)^18^ over 50 epochs. HPC specifications, training parameters, and performance metrics are given in ***Supplementary Tables S7*** and ***S8***.

***Supplementary Table S6*** *Modified U-Net architecture*

| **Block** | **Channels** | **Spatial size** |
| --- | --- | --- |
| Input | 1 | 224 × 224 |
| Conv1 | 64 | 112 × 112 |
| MaxPool | 64 | 56 × 56 |
| e1 | 256 | 56 × 56 |
| e2 | 512 | 28 × 28 |
| e3 | 1,024 | 14 × 14 |
| e4 | 2,048 | 7 × 7 |
| Central blocks | 1,024 | 7 × 7 |
| up4 | 512 | 14 × 14 |
| up3 | 256 | 28 × 28 |
| up2 | 128 | 56 × 56 |
| up1 | 64 | 112 × 112 |
| Output | 1 | 224 × 224 |

***Supplementary Table S7*** *U-Net training parameters and HPC specifications*

| **Parameter** | **Value** |
| --- | --- |
| GPU | 1 × NVIDIA A100 (80 GB) |
| CPU cores allocated | 8 |
| RAM allocated | 32 GB |
| Peak memory used | ~3.26 GB |
| Total wall-clock time | 6 h 10 min 51 s |
| Total CPU time | 16 h 08 min 07 s |
| Epochs | 500 |
| Input resolution | 224 × 224 px |
| Loss function | Dice + Weighted Cross-Entropy |
| Optimiser | AdamW |
| Learning rate scheduler | OneCycleLR |
| Mixed precision | Enabled |
| Software environment | PyTorch 2.1.2 (CUDA 12.1), torchvision 0.16.0, h5py 3.9.0, scikit-image 0.22.0, Python 3.11 (GCC 12.3.0) |

***Supplementary Table S8*** *U-Net internal performance metrics*

| **Metrics** | **Values** |
| --- | --- |
| Training accuracy | 93.76% |
| Best Dice score | 0.9366 |
| Best IoU score | 0.8242 |
| Final training loss | 0.1842 |
| Final validation loss | 0.2015 |

**Supplementary Note 5**

***Volumetric mesh generation***

Following slice-by-slice U-Net inference, 2D segmentation masks were stacked to reconstruct a 3D binary volume preserving the voxel spacing of the original CT scan, corresponding to the Mesh Generation and 3D Assembly stage (***Fig. 1****,* ***S2,*** *Stage 6*). The volume underwent three sequential refinement steps prior to surface extraction: region growing from high-confidence seed voxels (seed threshold = 0.5) outward to neighbouring voxels (grow threshold = 0.15)^19^; Otsu-scaled intensity filtering (scale = 0.60) to exclude non-bone attenuation values; and morphological operations (dilation and closing, radius = 1 voxel) to connect fragments and suppress residual noise. The marching cubes algorithm^20^ was then applied at an iso-level of 0.5 to extract a triangulated surface mesh. Mesh generation parameters are summarised in ***Supplementary Table S9***. Per-specimen mesh output quality metrics, including Dice, IoU, vertex counts, face counts, and generation runtimes, are reported in ***Table 1*** of the main text.

***Supplementary Table S9*** *Volumetric mesh generation parameters*

| **Parameter** | **Value** | **Purpose** |
| --- | --- | --- |
| Seed threshold | 0.50 | Minimum confidence to initialise bone region |
| Grow threshold | 0.15 | Lower bound for region expansion into adjacent voxels |
| Otsu scale | 0.60 | CT intensity gating to exclude non-bone regions |
| Marching cubes iso-level | 0.50 | Surface extraction threshold |
| Morphological radius | 1 voxel | Dilation and closing to connect fragments |

**Supplementary Note 6**

***Per-specimen registration phase decomposition: PPR, ICP, and C2M statistics for all six external validation specimens.***

Here we provide the granular, three-phase registration decomposition that underlies the assemblage-level results and tier classification reported in the main text (**Table 2**; **Fig. 4**). For each specimen, the Phase I Point Pair Registration (PPR)^21^ RMS, Phase II Iterative Closest Point (ICP) RMS, and Phase III Cloud-to-Mesh (C2M) signed distance statistics are tabulated alongside the corresponding voxel-normalised RMS Ratios, i.e., the raw registration outputs expressed in multiples of voxel size. The RMS Ratio is included to enable like-for-like comparison across specimens scanned at different physical resolutions (0.0154–0.0336 mm). The Phase I PPR was performed via manual landmarking to establish a common coordinate system; Phase II ICP was performed at 80% theoretical overlap for *Marmorerpeton wakei* Specimen 1, *Marmorerpeton wakei* Specimen 2, and Mammaliaformes indet. (specimens with significant non-overlapping geometry) and at 100% for the other three; in all six cases, Phase III C2M signed distances were computed on the full untrimmed point clouds (~50,000 points per specimen) regardless of the ICP overlap setting. A scale factor of 1.0 was held fixed across all phases for all specimens to prevent the algorithm from artificially expanding or contracting the AI mesh to minimise distances to noise clusters.

***Supplementary Table S10*** *‘Salamander A’ (Specimen 1, MorphoSource: 00084381)*

| **Voxel Size:** 0.015357 mm | | | | |
| --- | --- | --- | --- | --- |
| **Phase** | **Metric** | **Value (mm)** | **RMS Ratio** | **Interpretation** |
| I. PPR | RMS Error | 0.0896 | 5.83 | Good coarse alignment for high-resolution dataset. |
|  | Scale Factor | 1.0 (fixed) | N/A | No distortion in landmarking phase. |
| II. ICP  (100% overlap) | Final RMS | 0.1940 | 12.63 | Refinement affected by high-frequency surface noise. |
|  | Scale Factor | 1.0 (fixed) | N/A | Volumetric preservation confirmed at fixed scale. |
| III. C2M | Mean signed distance | 0.0632 | 4.11 | High global alignment accuracy (~4 voxels at 0.0154 mm). |
|  | Standard deviation | 0.1816 | 11.82 | Refined variance due to fine textural details. |
|  | Hausdorff (max) | ~10.469 | 681.6 | Outliers from distant fragmented noise plumes. |

***Supplementary Table S11*** *‘Salamander A’ (Specimen 2, rescan of MorphoSource: 000071513)*

| **Voxel Size:** 0.022798 mm | | | | |
| --- | --- | --- | --- | --- |
| **Phase** | **Metric** | **Value (mm)** | **RMS Ratio** | **Interpretation** |
| I. PPR | RMS Error | 0.0844 | 3.70 | Acceptable initial fit (>2 voxel size) |
|  | Scale Factor | 1.0 (fixed) | N/A | Baseline alignment establishes common coordinates |
| II. ICP (100% overlap) | Final RMS | 0.1417 | 6.21 | Robust alignment of core masses |
|  | Scale Factor | 1.0 (fixed) | N/A | Zero size difference, confirming strict geometric truth |
| III. C2M | Mean signed distance | −0.0170 | 0.75 | Sub-voxel accuracy; pipeline excludes matrix halo |
|  | Standard deviation | 0.1430 | 6.27 | Local surface variance dominates the residual signal |
|  | Hausdorff (max) | ~2.681 | 117.6 | Sparse outliers from binary thresholding noise |

***Supplementary Table S12*** Mammaliaformes indet. *(MorphoSource: 000693738)*

| **Voxel Size:** 0.024300 mm | | | | |
| --- | --- | --- | --- | --- |
| **Phase** | **Metric** | **Value (mm)** | **RMS Ratio** | **Interpretation** |
| I. PPR | RMS Error | 0.1641 | 6.75 | Stable initial fit for fragmented remains. |
|  | Scale Factor | 1.0 (fixed) | N/A | Rigid transformation baseline. |
| II. ICP (80% overlap) | Final RMS | 0.0289 | 1.19 | Best ICP fit in assemblage; near-perfect core alignment. |
|  | Scale Factor | 1.0 (fixed) | N/A | Precise geometric preservation. |
| III. C2M | Mean signed distance | 0.1543 | 6.35 | Accurate central alignment within ~6 voxels. |
|  | Standard deviation | 0.6376 | 26.24 | Spread inflated by ring-artefact banding (Avizo only). |
|  | Hausdorff (max) | ~4.894 | 201.39 | Reconstruction artefact noise correctly filtered. |

***Supplementary Table S13*** *Marmorerpeton wakei (Specimen 1, MorphoSource: 000700518)*

| **Voxel Size:** 0.033560 mm | | | | |
| --- | --- | --- | --- | --- |
| **Phase** | **Metric** | **Value (mm)** | **RMS Ratio** | **Interpretation** |
| I. PPR | RMS Error | 0.0812 | 2.42 | Lowest PPR ratio in assemblage; near-identity transform. |
|  | Scale Factor | 1.0 (fixed) | N/A | Global orientation established without warping. |
| II. ICP (80% overlap) | Final RMS | 0.1648 | 4.91 | Sub-micron rotations; translations <0.02 mm. |
|  | Scale Factor | 1.0 (fixed) | N/A | Confirms no artificial scaling. |
| III. C2M | Mean signed distance | 0.3840 | 11.44 | Elevation driven by partial overlap. |
|  | Standard deviation | 0.8130 | 24.22 | Highest spread in assemblage; non-matching peripheral fragments. |
|  | Hausdorff (max) | ~10.47 | 311.95 | Distant outlier fragments not representative of core fit. |

***Supplementary Table S14*** *Marmorerpeton wakei (Specimen 2, MorphoSource: 000700519)*

| **Voxel Size:** 0.032451 mm | | | | |
| --- | --- | --- | --- | --- |
| **Phase** | **Metric** | **Value (mm)** | **RMS Ratio** | **Interpretation** |
| I. PPR | RMS Error | 0.3634 | 11.20 | Challenging fit due to vertebral complexity. |
|  | Scale Factor | 1.0 (fixed) | N/A | Global orientation established. |
| II. ICP (80% overlap) | Final RMS | 0.1730 | 5.33 | Successful rigid refinement of complex mass. |
|  | Scale Factor | 1.0 (fixed) | N/A | Confirms no artificial warping occurred. |
| III. C2M | Mean signed distance | 0.3613 | 11.13 | Stable alignment despite high fragmentation. |
|  | Standard deviation | 0.6380 | 19.66 | Variance driven by matrix ghosting plumes. |
|  | Hausdorff (max) | ~5.455 | 168.10 | Lowest Hausdorff among 80%-overlap specimens. |

***Supplementary Table S15*** *Docodonta (undescribed, MorphoSource: 000721884)*

| **Voxel Size:** 0.025225 mm | | | | |
| --- | --- | --- | --- | --- |
| **Phase** | **Metric** | **Value (mm)** | **RMS Ratio** | **Interpretation** |
| I. PPR | RMS Error | 0.0902 | 3.57 | Strong coarse alignment of intricate bone mass. |
|  | Scale Factor | 1.0 (fixed) | N/A | Landmark-based registration baseline. |
| II. ICP (100% overlap) | Final RMS | 0.2421 | 9.60 | ICP convergence into local minima at fine bony processes. |
|  | Scale Factor | 1.0 (fixed) | N/A | Volumetric integrity preserved. |
| III. C2M | Mean signed distance | 0.0158 | 0.63 | Best mean ratio in assemblage; sub-voxel accuracy. |
|  | Standard deviation | 0.2351 | 9.32 | Surface chatter from scattered micro-debris. |
|  | Hausdorff (max) | ~4.253 | 168.61 | Noise outliers correctly rejected by AI pipeline |

***Methodological note on the ICP overlap parameter and fixed scale.***

The 80% theoretical overlap parameter was selected for the three specimens with significant non-overlapping geometry (multi-element vertebral complexes for both *Marmorerpeton wakei* specimens; severe ring artefacts and fragmentation for Mammaliaformes indet.). At this setting, ICP identifies and discards the worst-fitting 20% of point correspondences in each iteration, allowing the rigid alignment to converge on the core shared anatomy without being biased by outlier fragments. The Phase II ICP RMS values for these specimens are therefore trimmed statistics describing the fit of the shared region, and are not directly comparable to the 100%-overlap ICP RMS values of the other three specimens. The Phase III C2C signed distances, which were computed on the full untrimmed point clouds in all six cases, provide the consistent cross-specimen measurement basis and are the values used for tier classification in the main text. The fixed scale factor of 1.0 across all phases ensures that the registration metrics test geometric fidelity rather than scaled goodness-of-fit; this is essential when comparing AI segmentation against thresholding outputs, since allowing the algorithm to adjust scale would mathematically minimise distances by warping the AI generated cloud toward artefact noise clusters and produce a misleadingly low residual.

15	Ronneberger, O., Fischer, P. & Brox, T. in *International Conference on Medical image computing and computer-assisted intervention.* 234-241 (Springer).

16	Sudre, C. H., Li, W., Vercauteren, T., Ourselin, S. & Jorge Cardoso, M. in *International Workshop on Deep Learning in Medical Image Analysis.* 240-248 (Springer).

17	Loshchilov, I. & Hutter, F. Decoupled weight decay regularization. *arXiv preprint arXiv:1711.05101* (2017).

18	Smith, L. N. & Topin, N. in *Artificial intelligence and machine learning for multi-domain operations applications.* 369-386 (SPIE).

19	Adams, R. & Bischof, L. Seeded region growing. *IEEE Transactions on pattern analysis and machine intelligence* **16**, 641-647 (1994).

20	Lorensen, W. E. & Cline, H. E. in *Seminal graphics: pioneering efforts that shaped the field*	347-353 (1998).

21	Horn, B. K. Closed-form solution of absolute orientation using unit quaternions. *Journal of the optical society of America A* **4**, 629-642 (1987).
